# LeGO-Teknik reveals that the neurogenesis pathway is a clone-specific hallmark of brain metastasis in breast cancer

**DOI:** 10.1101/2025.07.23.666035

**Authors:** Jean Berthelet, Samuel C. Lee, Michael Roach, Sreeja R. Gadiaplly, Nora Liu, Farrah El-Saafin, Yunjian Wu, Caroline Bell, Sharon Li, Andres Vallejo-Pulido, Fernando J. Rossello, Emmanuelle Charafe-Jauffret, Christophe Ginestier, Matthias Ernst, Bhupinder Pal, Quentin Gouil, Jose Polo, Belinda Yeo, Luciano Martelotto, Delphine Merino

## Abstract

Breast cancer exhibits substantial inter- and intra-patient heterogeneity. Yet the molecular features underlying this diversity and their roles in tumour progression are not yet fully elucidated. Within a primary tumour, certain clones possess a unique capacity to metastasize to distant organs, such as the brain, and this propensity may be associated with specific gene expression profiles. To investigate this phenomenon, we developed a new optical barcoding library, *LeGO-Teknik*, compatible with high resolution imaging, flow cytometry, single-cell RNA sequencing and machine learning computational methods. Using this new tool, we systematically assessed the fitness and organ tropism of multiple human breast cancer clones in spontaneous metastasis assays. Linking transcriptomic profiles of individual cancer cells to their clonal identities and metastatic behaviours, we identified that clones predisposed to forming brain metastases exhibit distinct molecular signatures. Notably, individual cells from these clones showed enriched expression of neurogenesis-related genes. Using SCENIC transcription factor analysis, we found that some of these features are driven by the transcription factor SOX4. This hallmark, likely due to a process of cellular reprogramming, is a remarkable example of organ mimicry during clonal selection. Together, this work provides critical insights into brain metastasis and highlights the potential for identifying predictive biomarkers and therapeutic targets for metastatic breast cancer.

## Introduction

The prognosis of patients with breast cancer significantly hinges on whether malignant cells have spread to distant organs before diagnosis and therapeutic intervention ^1^. In particular, brain metastases, occurring in 10-15% of women with stage 4 breast cancer ^2^, are associated with the poorest prognosis ^3^ and are inherently unpredictable. Notably, patients with triple negative breast cancer (TNBC) and HER2-positive breast cancer have a higher likelihood of developing brain metastases as a first site of relapse compared to those with ER-positive disease ^4^. This suggests that the intrinsic properties of cancer cells may dictate organ-specific tropism. Within a given subtype, sequencing analyses of patient samples and cancer cell lines have revealed that tumour cells with a proclivity for brain metastasis exhibit distinct transcriptomic profiles ^5,6^. Furthermore, the brain represents a highly selective tumour microenvironment (TME) for the survival and growth of metastasis ^7^ and specific clones may have a selective advantage in colonising this organ ^8,9^. Investigating the level of inter- and intra-tumour heterogeneity associated with brain metastasis in a holistic way is a promising approach to identify the hallmarks of brain metastasis, and improve their prediction, prevention and treatment.

Cellular barcoding is a powerful tool for identifying and profiling individual clones associated with specific patterns of organ tropism ^10^. Lentiviral-based barcoding strategies have been employed to label cancer cells and track their progeny (also referred to as clones) as they colonise various organs, including brain metastases ^5,11^ and brain tumours ^12^. In particular, optical barcoding strategies using LEntiviral Gene Ontology (LeGO) vectors have proven successful for visualising the spatio-temporal distribution of the clones via imaging ^13–16^. This technology relies on fluorescent tags, stably integrated into the genomes of cancer cells, enabling their transmission to daughter cells. We previously demonstrated that combining five fluorescent LeGO tags (eBFP2, Sapphire, Venus, tdTomato and Katushka, referred to as BSVTK) can be used to track up to 31 barcoded cancer clones in various TMEs using flow cytometry and imaging ^17^. Additionally, the advantage of optical barcoding over other technologies, such as transcribed genetic barcoding library, is the non-destructive and real-time detection of the tags. This feature enables the easy isolation of all 31 clones via Fluorescence-activated Cell Sorting (FACS) for subsequent re-transplantation experiments and multi-omics analysis (this process of ‘clone-picking’ has been extensively discussed previously ^10^). This technology provides a robust platform for studying clonal selection and adaptation in xenograft models. However, a limitation of LeGO fluorescent tags is that they cannot be identified in single cell sequencing data, restricting the molecular characterisation of individual cells within and across barcoded clones.

To address this limitation, we developed an enhanced version of the LeGO vectors, termed LeGO-Teknik (or LeGOTek), which is compatible with 10X Genomics 3’ gene expression kits. Here, we demonstrated that this new library enables reliable identification of LeGO-Teknik tags in scRNAseq data. Using this technology, we investigated brain metastasis, linking the clonal identity of individual cells to their transcriptomic profile and fitness across different TMEs, including the brain. We also leveraged the multiplexing capability of this library to compare transcriptomic differences between clones and models at the single-cell resolution. Profiling of individual cancer cells, combined with their unambiguous labelling via the LeGO-Teknik barcodes, enabled us to apply machine learning and Single-Cell rEgulatory Network Interference and Clustering (SCENIC) analyses to identify genes and transcription factor programs that define brain tropism and tumour progression at the clonal and single-cell level. Our results reveal that cancer clones with a high proliferative potential in multiple metastatic sites, including the brain, are enriched in genes related to neuron differentiation and neurogenesis, genes typically expressed in healthy nervous system tissues.

## Results

### The new LeGO-Teknik library enables the detection of optical tags in single cell RNA sequencing

Optical barcoding of target cells is achieved using LeGO lentiviruses at high Multiplicity Of Infection (MOI), enabling the generation of a mixture of cells with multiple combinations of tags, identifiable by flow cytometry and imaging ^15^. Previously, we demonstrated that cancer cells, including MDA-MB-231, can be tagged with eBFP2, Sapphire, Venus, tdTomato and Katushka (BSVTK) fluorescent barcodes ^17^. The resulting combination of 31 colours (Extended Data Fig. 1a; colours maintained throughout the manuscript) can be distinguished by flow cytometry and confocal imaging. However, while individual clones can be FACS-sorted and subsequently analysed for bulk or single-cell sequencing (one clone per single cell capture) ^17^, the transcribed fluorescent tags were not reliably detected in scRNA-seq data (Extended Data Fig. 1b-e), impairing the ability to study the diversity of a mixed cell population (e.g. several clones per single cell capture). This limitation arises due to a shared 1,300 bp sequence containing the Woodchuck hepatitis virus Post-transcriptional Regulatory Element (WPRE) in the 3’ untranslated region (3’UTR) of the LeGO sequence. Indeed, when using the 10X Genomics 3’ single-cell RNA chemistry to sequence a clone simultaneously expressing three fluorescent LeGO tags, eBFP2, Venus and tdTomato (Extended Data Fig. 1b), the 3’UTR upstream of the polyA signal could be detected in all cells (Extended Data Fig. 1c, right panel). However, sequencing read coverage for the coding sequence of the fluorescent tags was low, allowing for the reliable detection of only a single tag in a very small proportion of cells (Extended Data Fig. 1d, e). This issue compromises the clonal identity calling in scRNA-seq, as the presence or absence of all fluorescent tags must be identified to accurately determine the colour combination expressed in individual cells. Indeed, only ∼1.2% of cells were identified with the correct combination of fluorescent tags in this clone (Extended Data Fig. 1e).

To address this technical challenge, we developed a new collection of LeGO plasmids compatible with 10X Genomics’ 3’ single-cell sequencing (Fig. 1a). To ensure maximal reliability in tag detection, the sequencing library of these new vectors was designed to distinguish highly similar fluorescent protein sequences and maintain fluorescence levels comparable to conventional LeGO vectors. To achieve this, we retained the WPRE signal and introduced a ‘Teknik cassette’ near the PolyA sequence of the conventional LeGO vectors (Fig. 1b). This cassette contains a 30-nt barcode (unique to each fluorescent tag) and a sequence complementary to 10X Genomics 3’ Capture Sequence 2. By design, the Teknik feature library can be prepared separately from the gene expression (GEX) library containing the transcriptome of individual cells. This design allows both libraries to be indexed separately and sequenced together or independently, providing control over the sequencing depth of each experiment.

**Figure 1:**
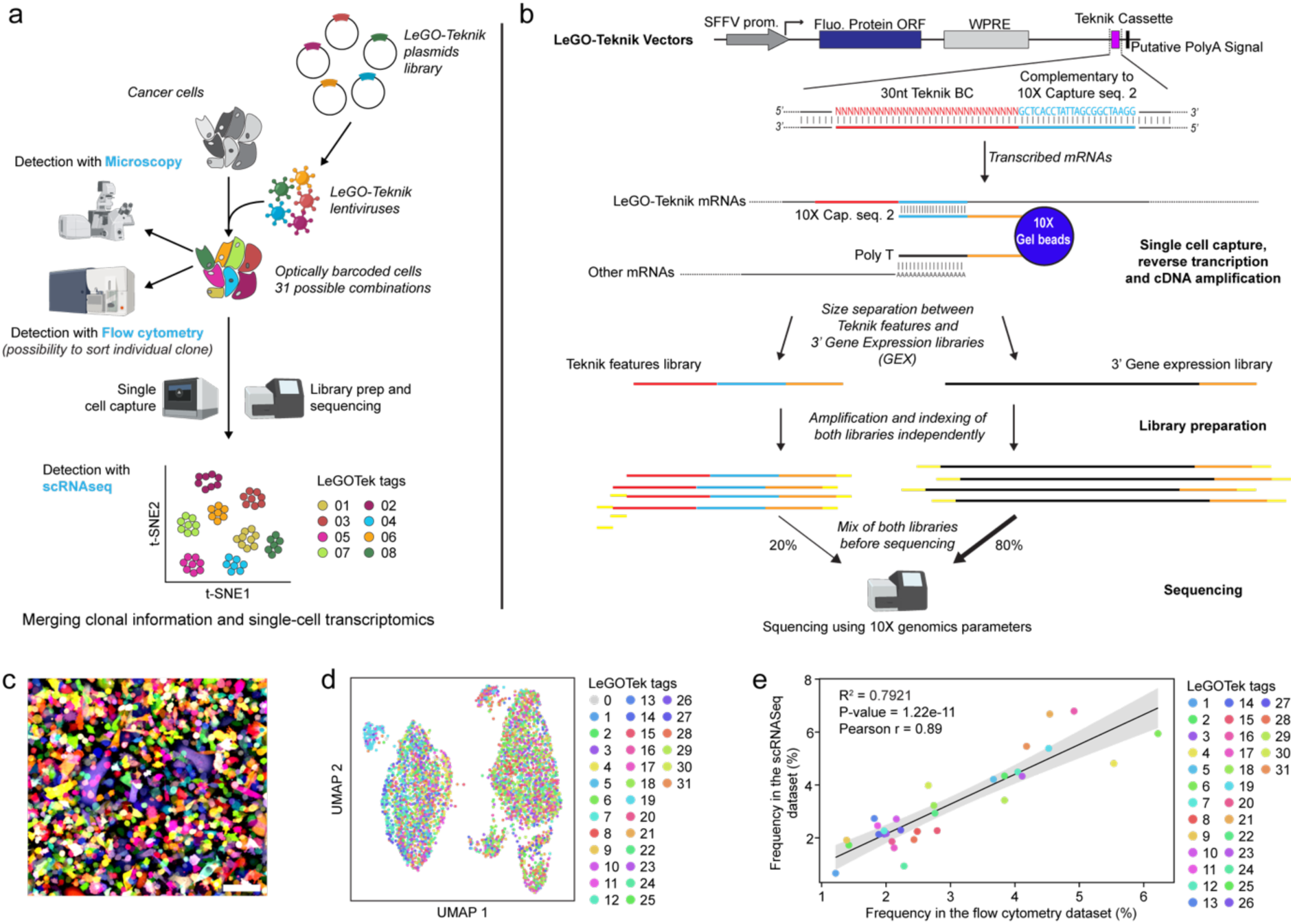
Design and validation of the new LeGO-Teknik optical barcoding library. **a**) LeGO-Teknik workflow overview. **b**) LeGO-Teknik vector design and library preparation workflow. **c**) Fluorescence imaging of MDA-MB-231 cells infected with LeGO-Teknik (scale bar = 100µm). **d**) UMAP projection of scRNA sequencing of MDA-MB-231 cells infected with LeGO-Teknik vectors, coloured by computationally decoded LeGO-Teknik tags. **e)** Correlation of the frequency of individual clones, represented by different colours as indicated in the legend, in flow cytometry (using fluorescence) and single cell RNA sequencing (using *in silico*-derived LeGOTek tag counts).

To validate this library, we infected MDA-MB-231 cells with a mix of BSVTK LeGO-Teknik viruses (Fig. 1c). We profiled ∼10,000 cells and found that the five optical tags could be detected in the feature library (Extended Data Fig. 1f-h), and the 31 colours could be computationally assigned to individual cells from the GEX library (Fig. 1d). Furthermore, the ratio of the 31 colours identified by scRNA-seq correlated with the ratio detected by flow cytometry (Fig. 1e), and the presence of LeGO-Teknik vectors did not affect the quality of the GEX dataset (Extended Data Fig. 1i-l). Together, these results demonstrate that this new library enables robust detection of optical tags in scRNA-seq, linking the transcriptome of individual cells to their clonal identity.

### LeGO-Teknik can be used to link clonal identity, transcriptome and behaviour

To investigate the hallmarks of breast cancer clones associated with specific clonal behaviour (i.e., metastatic fate and clonal fitness), we generated single-cell-derived clones from the MDA-MB-231 cell line labelled with BSVTK LeGO-Teknik. Individual colour-coded cells were isolated using FACS and amplified independently. From these cells, 13 clones with different colour combinations grew and their purity was confirmed by flow cytometry before pooling them in equal quantities (Extended Data Fig. 2a). A fraction of these cells was scRNA-seq profiled, while others were transplanted into the mammary fat pad of multiple recipient NSG mice (Fig. 2a). Single-cell gene expression analysis revealed that cells cultured *in vitro* clustered based on their colour (and therefore, clonal origin), demonstrating the presence of inter-clonal transcriptomic heterogeneity (Fig. 2b, Extended Data Fig. 2b-g).

**Figure 2:**
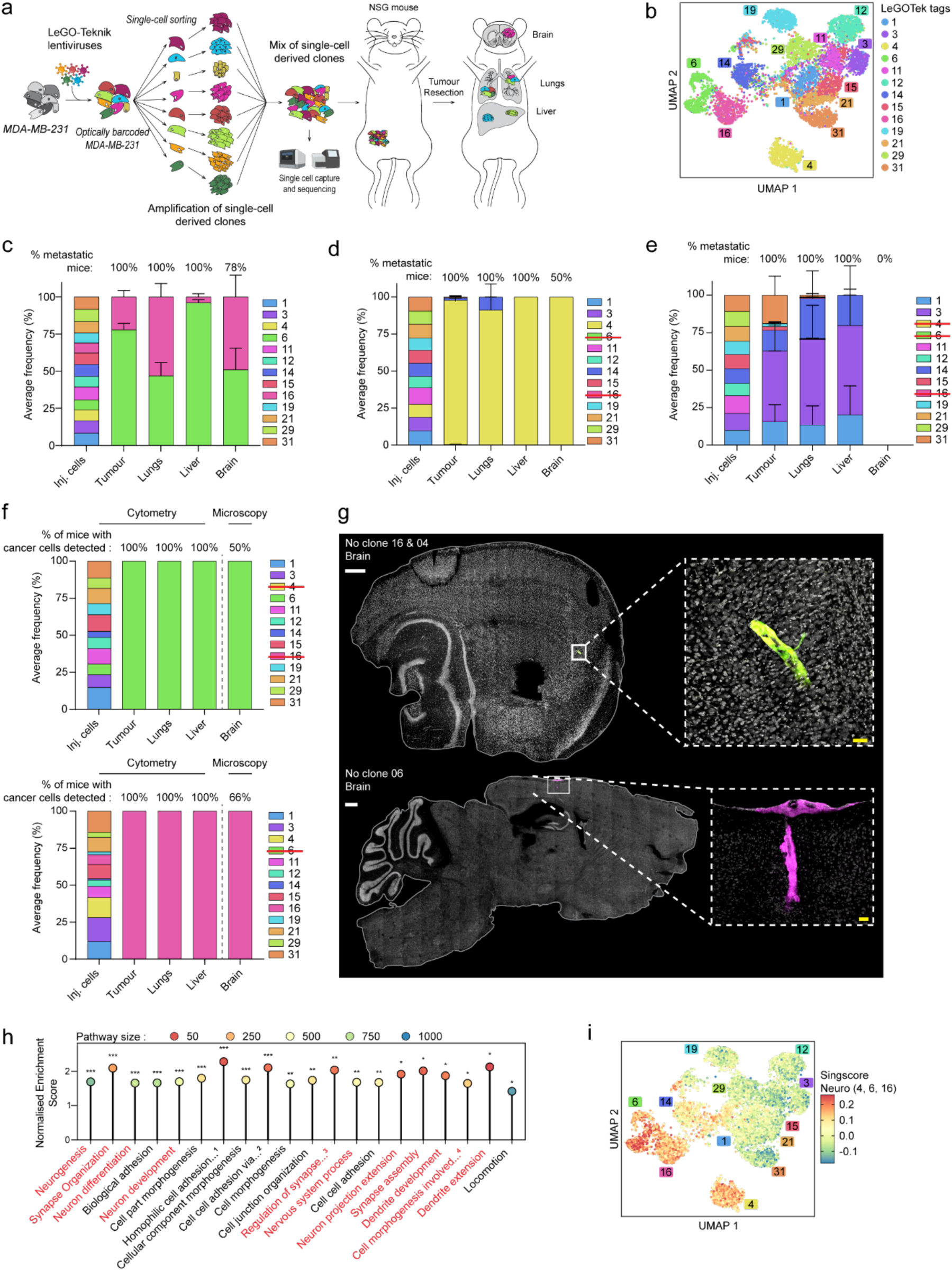
Leveraging LeGO-Teknik to link clonal identity with transcriptomic profiling and metastatic behaviour. **a**) Overview of the in vitro amplification of single-cell derived MDA-MB-231 clones labelled with unique LeGOTek tags, and their use in single cell sequencing and in vivo experiment. **b**) Single cell gene expression UMAP embeddings of LeGO-Teknik tagged single-cell derived MDA-MB-231 clone mix and coloured by their LeGOTek tags detection. **c–e**) Average frequency of the LeGOTek tags detected in the organs of mice harvested at metastatic ethical endpoint, after the tumour resection. The LeGOTek tags were detected and quantified using flow cytometry. The injected populations contained: all 13 LeGO-Teknik tagged clones (*c ; n=9 mice*), all clones except for clones 6 and 16 (*d ; n=6 mice*), and all clones except for clones 4, 6, and 16 (*e ; n=5 mice*). The percentage of mice where fluorescent cells were detected in specific organs is indicated above the bar graph. Error bars represent the standard-error of the mean. **f**) Average frequency of the LeGOTek tags detected in the organs of mice harvested at ethical endpoint after the tumour resection. The LeGOTek tags were analysed using flow cytometry in cell suspensions from the tumour, lungs and liver. The presence of barcode in the brain was analysed using confocal microscopy. The injected population comprised: all clones without clones 4 and 16 (n=6 mice, top) or all clones without clone 6 (n= 6 mice, bottom). The percentage of mice with fluorescent cells were detected in distinct organs is indicated above the bar graph. **g**) Brain metastasis confocal microscopy image from mouse injected without clone 16 and 4 (top) or without clone 6 (bottom) (white scale bar = 500 µm, yellow scale bar = 50 µm). **h**) Gene Ontology Biological Process pathway enrichment for differentially expressed genes between clones 4, 6, and 16 and the others. Pathways related to the nervous system are indicated in red. Significance is indicated (* p-value < 0.05; ** p-value < 0.01; *** p-value < 0.001). Abbreviations: 1, Homophilic cell adhesion via plasma membrane adhesion molecules; 2, Cell-cell adhesion via plasma membrane adhesion molecules; 3, Regulation of synapse structure or activity; 4, Cell morphogenesis involved in neuron differentiation. **i**) UMAP projection of the 13 LeGO-Teknik tagged clones coloured by Singscore scores for the differentially expressed genes between clones 4, 6, and 16 and others, that are part of the neurogenesis gene ontology biological process.

To test whether these clones, all derived from the same cell line, were genetically heterogeneous, DNA sequencing of 5 clones (clones 3, 4, 6, 16 and 31) was performed using Nanopore sequencing. Single nucleotide polymorphism (SNP) analysis revealed that these 5 clones were genetically distinct (Extended Fig. 2h), although some clones were more closely related to others (Extended Fig. 2i), in agreement with the scRNA-seq analysis (Fig. 2b).

We next assessed the ability of the 13 barcoded clones to grow in primary and metastatic sites. Following intra-mammary fat pad injection of all clones, primary tumours were resected, and mice were euthanised at metastatic ethical endpoint. The clonal composition of the primary tumours and metastatic organs was assessed by flow cytometry. Interestingly, despite the 13 clones being equally represented in the injected population, two clones (clones 6 and 16) dominated in both primary tumours and metastases (Fig. 2c, Extended Data Fig. 3a and b). This clonal dominance was observed consistently across multiple mice (Extended Data Fig. 3a), suggesting an intrinsic, non-random mechanism of metastatic spread. To assess the ability of the other clones to grow at primary and metastatic sites, we injected a pool of clones that lacked clones 6 and 16. Interestingly, the analysis of primary tumours revealed the emergence of new, hitherto under-represented clones (i.e. clone 4, and to a lesser extent clone 14) (Fig. 2d). While the tumours grew slower and mice survived longer until metastatic endpoint (Extended Data Fig. 3c, d), all mice still developed lung and liver metastases and 50% developed brain metastases (Fig 2d, Extended Data Fig. 3e). In this cohort, clone 4 was seen to be dominant across primary tumours and most metastases, and was the only clone detected in the brain (Fig. 2d and Extended Data Fig. 3e). We then repeated this experiment, this time excluding clones 4, 6, and 16. In this case, tumour development was further delayed and the mice survived longer compared to both the previously injected pools (Extended Data Fig. 3c, d). However, while all mice still developed lung and liver metastases, no cancer cells were detected in the brains of these mice (Fig. 2e, Extended Data Fig. 3f). Altogether, these results suggested that clones 4, 6 and 16 had a unique capacity to colonise the brain compared to the other clones.

To assess any potential cooperativity between clones with brain tropism, we injected either all clones except 4 and 16, or all clones except 6. The analysis revealed that in absence of clones 4 and 16, clone 6 dominated in both primary tumours and brain metastases. Similarly, in absence of clone 6, clone 16 was dominant (Fig. 2f, g). These results indicate that these clones can metastasise to the brain independently of one another. Of note, brain metastases obtained with this spontaneous model were micro-metastases, detected thanks to the high-resolution of LeGO tags using flow cytometry (Fig. 2c-e) and imaging (Fig. 2f, g), as these mice had to be euthanised due to large metastases in other organs (Fig. 2c-f).

To explore the molecular signatures associated with brain tropism, we compared the single-cell gene expression profiles of these three clones (4, 6, 16) to other clones (Fig. 2b). We identified 349 differentially expressed genes (Table 1). Interestingly, 6 of the top 10 genes enriched in a previously described MDA-MB-231-BrM variant generated by experimental metastasis assays and re-transplantations of brain metastases ^18^, were also upregulated in clones 16, 6 and 4 (MMP1, ROBO1, CST1, PCDH7, MAN1A1, AGR2). Our experiments, using spontaneous metastatic assays and cellular barcoding, suggest that these genes were likely to be already expressed in the parental line, rather than the result of cellular adaptation in the brain.

Focussing on pathway analysis, we identified over 40 enriched Gene Ontology (GO) Biological Process (BP) pathways (Table 2). Strikingly, this analysis revealed major classes of processes related to adhesion, locomotion, morphogenesis and several neurogenesis/neuron-associated pathways (Fig. 2h, in red, Table 2). When scoring individual cells for all differentially expressed genes (Extended data Fig. 4a) or exclusively for genes within the neurogenesis signature (Fig. 2i), we observed that the neurogenesis pathway was a clonal feature of clones 4, 6, and 16, as it was homogeneously expressed in individual cells from these clones. Furthermore, clone 4 and, to a lesser extent, clone 14 exhibited an intermediate phenotype compared to clones 6 and 16.

To determine whether the biased enrichment for the neurogenesis pathway was maintained in the primary tumour, we compared the transcriptomic profiles of clones 6 and 16 in *in vitro* cultures and in primary tumours using bulk RNA sequencing, using the LeGO-Teknik ability to sort individual clones based on their colour (Extended Data Fig 4b). Clone 3 was included as an example of a clone not found in brain metastasis. The results confirmed that the neurogenesis score identified *in vitro* was also present in clones 6 and 16 *in vivo* within the primary tumour (Extended Data Fig 4b). It also suggested that this gene signature did not completely rely on the presence of the tumour microenvironment.

In summary, these findings indicate a clonal nature of the MDA-MB-231 cell line with some clones endowed with a transcriptional profile including neurogenesis-related genes, associated with a competitive advantage to dominate at primary and metastatic sites, including the brain.

### The neurogenesis pathway is enriched in primary tumours from PDXs with brain tropism

We leveraged the multiplexing capacity of LeGO-Teknik to investigate the association between brain tropism and neurogenesis pathway in five drug-naive patient-derived xenografts (PDXs): one *ERBB2* amplified breast cancer (PDX-CRCM226, shortened to PDX-226) and 4 triple-negative breast cancer (TNBC) models (PDX-CRCM434 or PDX-434, PDX-CRCM412 or PDX-412, PDX-1432 and PDX-0066). Each PDX was labelled with one of the five LeGO-Teknik tags (Fig. 3a, Extended Data Fig. 5a). Notably, the PDXs were derived from primary tumours, except PDX-0066, which originated from a malignant pleural effusion. We then analysed the transcriptome of these PDXs after pooling cells from all five models in equal proportion. Barcode identities, representing PDX origins, were retrieved from the feature library, as described earlier. As expected, cancer cells clustered based on their barcode (i.e., patient origin) (Fig. 3b, Extended Data Fig. 5b-d). To assess the aggressiveness of these models in primary and metastatic sites, we injected a mixed population of cells from all five PDXs into the mammary fat pad of recipient mice. Although equal number of cells from each PDX were injected, PDX-434 and PDX-1432 demonstrated a clear competitive advantage at the primary site (Fig. 3c, d). However, only PDX-434 cells were detected in the brain of these mice (Fig. 3d, e). Consistent with these findings, PDX-434 and PDX-1432 were the fastest to generate primary tumours when injected individually (Extended Data Fig. 5a). However, in this context, all PDXs, except PDX-1432, demonstrated the ability to metastasize to the brain (Fig. 3f). When comparing the transcriptome of PDXs capable of forming brain metastases (all except PDX-1432) with the one that did not (PDX-1432), we observed that the neurogenesis pathway was among the top enriched pathways (Fig. 3g, Tables 3, 4). The overall ability of the PDXs to form brain metastases correlated with their neurogenesis scores, with PDX-434 displaying the highest score (Fig. 3h).

**Figure 3:**
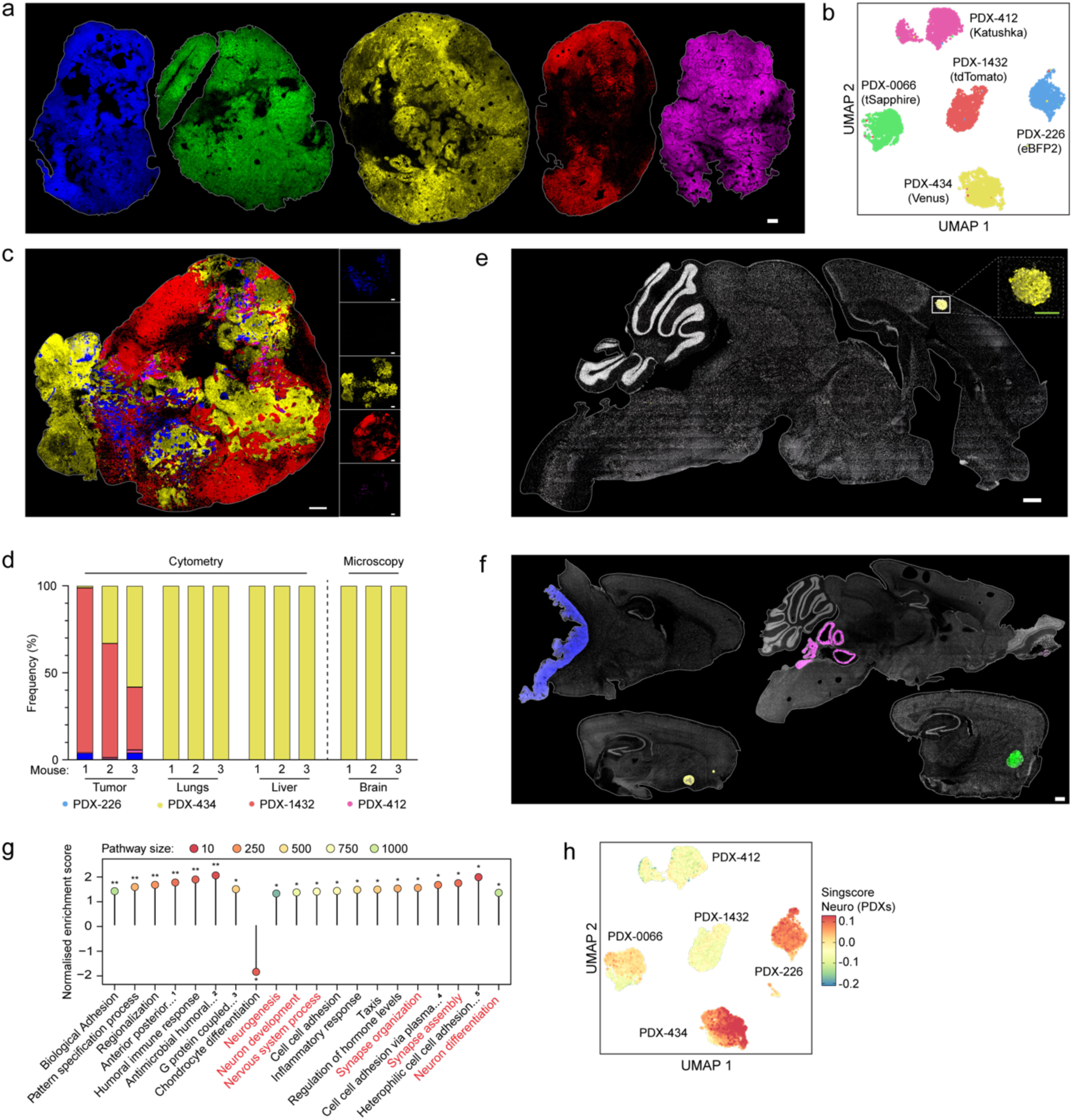
Metastatic potential of LeGO-Teknik tagged 5 Patient-Derived Xenografts. **a**) Confocal microscopy images of the primary tumours of 5 PDXs labelled with individual LeGOTek tag: PDX-226 with Teknik-eBFP2 (in blue), PDX-0066 with Teknik-Sapphire (green), PDX-434 with Teknik-Venus (yellow), PDX-1432 with Teknik-tdTomato (red) and PDX-412 with Teknik-Katushka (magenta) (scale bar = 500µm). **b**) Single cell gene expression UMAP embeddings of the 5 PDXs primary tumour captured and sequenced together. Each dot is coloured by computationally identified LeGOTek tags. **c**) Representative image of a primary tumour resulting from the injection of the 5 PDXs together (scale bar = 500µm). **d**) Frequency of each PDX detected in the organs of mice (*n=3 mice*). Primary tumours were harvested and analysed at the time of resection. Lungs, liver and brain were analysed at ethical metastatic endpoint. The LeGOTek tags were analysed using flow cytometry for the tumour, lungs and liver while the brain was analysed using confocal microscopy. The primary tumour was resulting from the injection of the mix of 5 PDXs. **e**) Representative brain metastasis confocal microscopy image from a mouse injected with the mix of 5 PDXs. Only PDX-434 (LeGOTek-Venus, yellow) was detected in the brain of these animals (white scale bar = 500µm, green scale bar = 200 µm). **f**) Representative image of brain metastasis by confocal microscopy, from a mouse injected with PDX-226 (LeGOTek-eBFP2, blue, **top left**), PDX-434 (LeGOTek-Venus, yellow, **bottom left**), PDX-412 (LeGOTek-Katushka, magenta, **top right**) and PDX-0066 (LeGOTek-Sapphire, green, **bottom right**) (scale bar = 500µm). Brain metastases were not detected in mice injected with PDX-1432 (LeGOTek-tdTomato, red). **g**) Gene Ontology Biological Process pathway enrichment of the differentially expressed genes between PDXs able to form brain metastasis (PDX-434, −0066, −412 −226) and other (PDX-1432). Nervous system-related pathways are indicated in red. Significance is indicated (* p-value < 0.05; ** p-value < 0.01; *** p-value < 0.001). **h**) UMAP projection of the scRNA sequencing analysis of the 5 PDXs, coloured by Singscore score of the differentially expressed genes (when comparing PDX-434, −0066, −412 −226 vs PDX-1432) that are part of the neurogenesis gene ontology biological process. Abbreviations for panel g: 1, Anterior-posterior pattern specification; 2, Antimicrobial humoral response; 3, G protein-coupled receptor signalling pathway; 4, Cell-cell adhesion via plasma membrane adhesion molecules; 5, Heterophilic cell-cell adhesion via plasma membrane cell adhesion molecules.

### Breast to brain metastasis is a clonally imprinted mechanism of organ mimicry

Experiments in multiple hosts with MDA-MB-231 cells demonstrate that the ability of cancer cells to metastasise to the brain is intrinsically programmed and suggest that the expression of neuronal genes might be, at least in part, associated with this process in this cell line and these PDXs. To further investigate the underlying mechanisms, we compared the signatures of genes deregulated in MDA-MB-231 brain metastatic clones (Table 1) to genes expressed in autopsy samples from healthy human tissues (GTEx). We found that expression of these genes was enriched in nerves, brain, and the pituitary gland compared to other organs (Fig. 4a). Notably, these genes are expressed in various cell types of the nervous system lineage, including neurons (Extended Data Fig. 6a).

**Figure 4:**
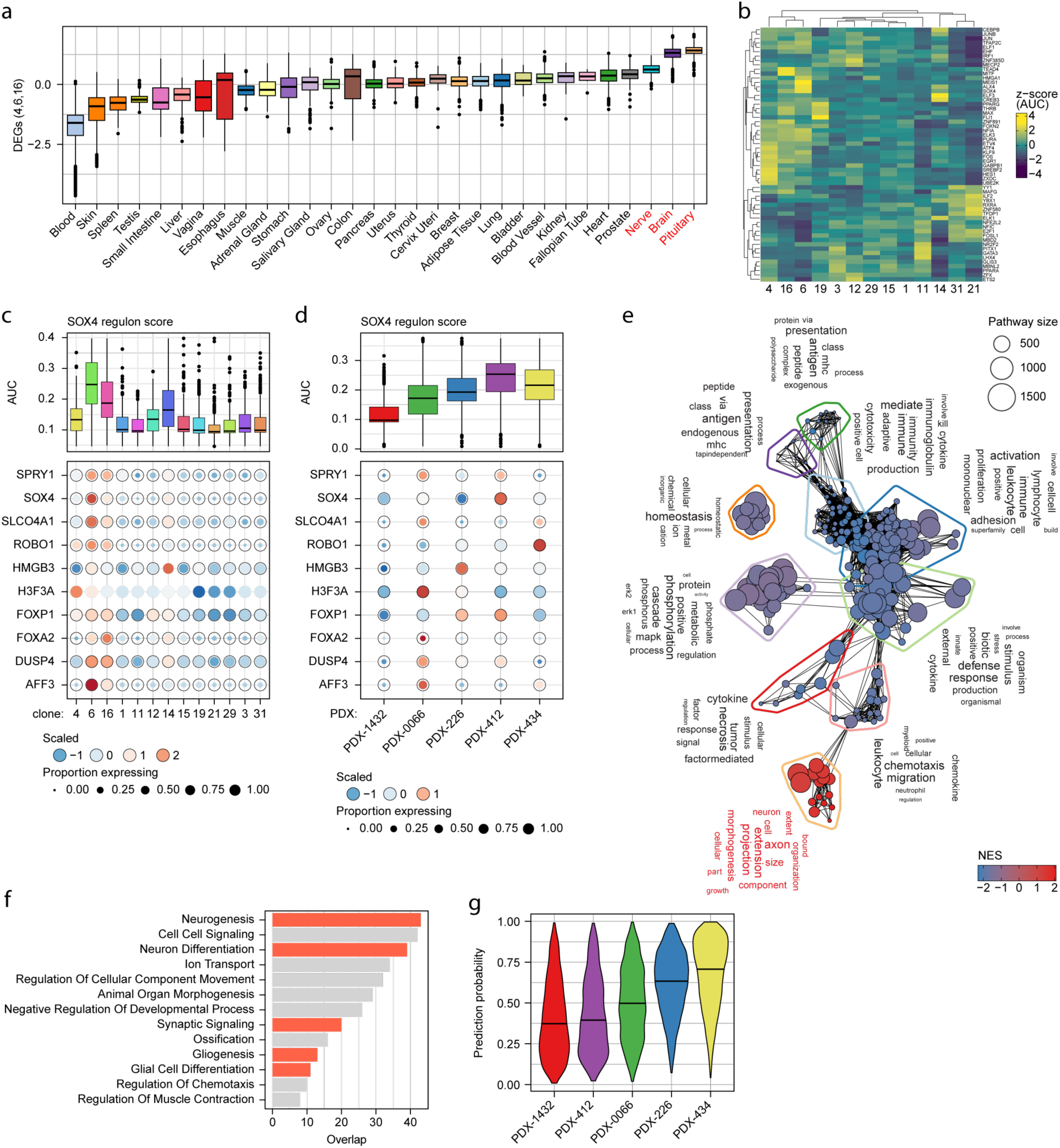
Gene signatures predictive of brain metastasis in the MDA-MB-231 cell lines and PDXs. **a**) Singscore scores (z-score transformed) of the MDA-MB-231 4/6/16 DE gene signature on human autopsy tissue samples from GTEx. **b**) SCENIC AUCell scores (z-score transformed) of pseudobulked clones for the top 5 marker regulons per clone. **c**) The high-confidence SOX4 regulon from SCENIC with per-clone AUCs (top panel) and individual gene expression scaled per gene (bottom panel). **d**) AUCell scoring of the SCENIC SOX4 regulon in the PDX models. **e**) vissE network of pathways with significant up (or down) regulation in SOX4 overexpressing clone 3 cells compared to control clone 3, textual themes for pathway modules (circled) are shown in word clouds. **f**) Pathway enrichment analysis of the 321 selected model genes, pathways with an FDR < 0.1 are displayed, with neural related pathways coloured in red. **g**) Validation of elastic net model performance in the PDX dataset, violin plots of the positive class prediction probabilities per PDX model.

We next explored whether the expression of neurogenesis-related signatures in specific MDA-MB-231 clones might result from a process of reprogramming through changes in their underlying regulatory networks ^19^. To do so, we used SCENIC, a method for single-cell regulatory network inference and clustering, designed to identify active transcription factors and their targets, or regulons ^20^. Of the 171 high-confidence regulons identified in the MDA-MB-231 data, a number showed marked inter-clonal heterogeneity (Fig. 4b, Table 5). Comparison between the brain metastatic competent and incompetent clones highlighted 46 regulons with statistically significant differences in their activity (Extended Data Figure 6b). Interestingly, we found that many of these regulons were also deregulated in PDXs (Extended data Fig. 6c). Among these, we found that the SOX4 regulon, previously shown to promote neuronal and neural progenitor differentiation ^21^, was significantly upregulated in both MDA-MB-231 clones and PDXs with brain tropism (Fig. 4c, d). Here the advantage of LeGO-Teknik over conventional LeGO is that it enabled gene expression profiling of individual cells in each clone or model, and visualisation of both the percentage of cells expressing each gene and their expression levels (Fig. 4c and d). These differences in expression levels and frequency illustrate the extent of heterogeneity within and between tumours, even for a given regulon.

To test whether SOX4 upregulation plays a role in the reprogramming of these cells, we overexpressed SOX4 in clone 3, showing a lower SOX4 regulon score and brain tropism compared to clones 4, 6 and 16 (Fig. 2e, 4c, Extended Data Fig. 6d). RNA sequencing analysis revealed widespread transcriptomic changes to clone 3 after over-expression of SOX4 (1,694 upregulated and 2,034 downregulated genes at an FDR < 0.05) with top deregulated genes including CD24, DMP1, DRAXIN, and CXCL8 (Extended Data Fig. 6e, Table 6). Reprogramming towards a clone 6-like state was confirmed by scoring of wild-type clone 6 and 3 cells with the SOX4 overexpression signature (TREAT test selected genes: 150 up, 150 down), where clone 6 cells had positive scores (like clone 3-SOX4) and clone 3 negative (Extended Data Fig. 6f). Pathway analysis of the SOX4 overexpressing cells highlighted two distinct sets of pathway modules using vissE: a set of down-regulated modules mostly associated with the innate immune response and a module of upregulated pathways associated with neuronal function and in particular axonogenesis (Fig. 4e, Extended Data Fig. 6g, Table 7).

Collectively, these findings suggest that clones predisposed to metastasise to the brain may have undergone a process of reprogramming to activate gene expression programs typically associated with a neuronal cell identity, through transcription factors such as SOX4.

### Leveraging the use of machine learning classification to identify a predictive signature of brain tropism

A distinctive feature of the LeGO-Teknik approach is its ability to decouple cell annotation; specifically, clonal identity, from transcriptome profiling. This contrasts with non-barcoded single-cell data, where cell type labels are typically assigned through gene expression analysis. Consequently, the LeGO-Teknik approach enables the use of machine learning to identify relevant transcriptomic features associated with brain tropism without the risk of overfitting by leakage between the response and predictor variables. To capitalise on this advantage, we developed a supervised learning classifier to predict the metastatic potential of individual cells using single cell gene expression data. An elastic net classifier was trained to distinguish MDA-MB-231 clones with brain metastatic potential (clones 4, 6, and 16; positive class) from non-brain metastatic clones (clones 1, 3, 11, 12,14, 15, 19, 21, 29, and 31; negative class). Using 80% of the cells as the training split, we obtained a final model containing 321 genes with non-zero weights (Extended Data Fig. 7a). The model’s validity was confirmed using the hold-out 20% testing split, where it classified cells with 96% accuracy (Extended Data Fig. 7b, Tables 8 and 9). An alternative training scheme was considered, where leave-one-out testing was done per clone. Using this approach, removing either clones 4 or 14 from the training dataset showed reduced testing accuracy but all other clones were still classified with high accuracy (Table 10). As clone 4 is less aggressive in the primary tumour than clones 6 and 16, this suggests that the model trained using clone 4 as held out test data was likely confounding clonal aggression and brain metastatic competence in the positive class of the training data. As an orthogonal validation of the model, single-sample scoring of the 321 selected genes on bulk MDA-MB-231 gene expression data revealed enrichment of clones 6 and 16 compared with clone 3 (Extended Data Fig. 6c). Additionally, gene set testing confirmed the enrichment of neural development-related pathways, amongst others (Table 8 and 11, 43 out of 321 genes, Figure 4f).

To evaluate whether this could be generalised beyond the MDA-MB-231 context, we applied this model to score the brain metastatic potential of single cells from barcoded PDX samples. Probabilities of cells being assigned to the positive class were estimated. The highly aggressive and brain-metastatic PDX-434 displayed higher scores, while the aggressive but non-brain metastatic PDX-1432 showed low scores (Fig. 4g, Extended Data Fig. 6d, e). Interestingly, the remaining three PDXs, which were less aggressive but capable of brain metastasis, showed intermediate scores. These results underscore a shared gene expression signature of brain metastatic competence between both clonally defined MDA-MB-231 populations and PDX lines from multiple patients.

While the overall brain metastasis signature, which included genes from neurogenesis-related pathways, could predict brain tropism, other genes and pathways related to morphogenesis and locomotion were also included in this model (Tables 8 and 11, Fig. 4f). These results suggest that brain metastases might be the result of the integration of multiple biological pathways and physiological functions, involved in both aggressiveness and brain tropism.

## Discussion

The spread of breast cancer to specific organs remains unpredictable. Using the molecular characteristics of primary tumours to predict the likelihood and site of relapse is an appealing strategy for the management of breast cancer patients. To address this clinically relevant challenge, we developed a new optical barcoding library, LeGO-Teknik. This optical barcoding library is compatible with multiomic analysis, including 3D imaging, flow cytometry, and scRNAseq, and enables the quantification, visualisation and characterisation of cancer clones at single-cell resolution. Indeed, as with conventional LeGO vectors, LeGO-Teknik tags can be used not only to study the fate and fitness of individual clones by imaging or flow cytometry, but also to isolate individual clones by FACS. This unique ‘non-destructive’ feature of clonal tracking enables clonal characterisation of live cells, followed by any type of analysis. In addition, the delineation of barcode identity in scRNAseq via Feature sequencing allowed us to link metastatic behaviour detectable by high-resolution imaging (e.g. brain metastasis) to the transcriptomic profile of the cells, for each and every clone. Indeed, this library enables the multiplexing of up to 31 populations in one 10x sequencing capture, lowering the cost of single-cell RNA sequencing experiments and limiting batch effects. The compatibility of optical barcoding with flow cytometry also allows for the normalisation of individual populations by FACS prior to single-cell capture, for instance, to enrich the proportion of a minor clone, which can’t be easily achieved for 31 clones by genetic barcoding. Finally, we demonstrated that this technology is compatible with machine learning classification and single-cell analytic tools, such as SCENIC, offering new avenues to identify signatures associated with clonal behaviour, at the single-cell level.

Using a unique collection of human xenograft models spontaneously metastasising to the brain, including the MDA-MB-231 cell line and 4 drug naive PDXs, we uncovered the hallmark of clones and models with a higher likelihood to spread to the brain. In particular, we discovered that clones likely to metastasise to the brain express gene programs normally associated with neural cells, a possible mechanism of neural mimicry predisposing cells to survive and grow in the brain. These results are in agreement with previous studies showing that TNBC cells expressing neuronal related genes are likely to spread to the brain ^22^ and that neuronal activity promotes invasion and metastases in the brain TME ^23,24^. Furthermore, a previous analysis of non-treated brain and non-brain metastases in patients with melanoma showed that brain metastases were enriched in genes involved in neural differentiation as well as adhesion and morphogenesis ^6^. Similar neuronal features were described in lung cancer^25^. The expression of these genes, usually expressed in neural cells, are likely to confer a selective advantage once the cells reach the brain. However, our results demonstrate that these genes are deregulated at the clonal level in MDA-MB-231 cells, and that this process may occur before cancer cells reach the brain. Furthermore, we confirmed in several PDXs derived from drug naïve primary tumours, that models with brain tropism upregulate genes from the neurogenesis pathway prior to metastatic spread.

The stimuli and mechanisms involved in the upregulation of this pathway are still unclear. However, our results in the MDA-MB-231 cell line revealed that clones able to metastasise to the brain upregulate targets of several neuronal differentiation transcription factors. Interestingly, one of them, SOX4 is associated with metastasis and poor prognosis in breast cancers from other types ^26–28^. Furthermore, SOX4 was previously shown to be upregulated in an MDA-MB-231-brain metastatic variant compared to the parental cell line ^18^. Here, we demonstrated that upregulation of SOX4 in cancer cells results in an enrichment of genes associated with neural differentiation and axonogenesis, that is likely to contribute to brain metastasis. While the role of SOX4 in regulating immunogenicity in cancers (including TNBC) has been well characterised ^28^, the induction of a pro-neural/axonal program by SOX4 has been understudied in the context of breast cancer. However, this result is consistent with a similar phenotype observed in lung cancer where SOX4 knockout was seen to deregulate genes associated with a neural-like phenotype ^29^. Interestingly, axon-like protrusions and neuronal reprogramming have been identified in small-cell lung cancers, and have been described as promoting metastasis in several organs, including the brain ^30^.

While it will be important to determine if SOX4 overexpression, alone, is sufficient to increase brain tropism in specific clones or models, several transcription factors and pathways are likely to be required. Indeed, SCENIC analysis identified other regulons upregulated in clones 4, 6 and 16, that involved in pro-neural functions or synaptic plasticity such as NFIA and EGR1, respectively ^31–33^. Indeed, our study shed light on the heterogeneity of mechanisms involved, even between clones from the MDA-MB-231 cell lines, and highlighted that one pathway, and one gene, alone, is unlikely to be predictive of brain metastasis. Indeed, in addition to the neurogenesis pathway, other pathways such as locomotion, morphogenesis and adhesion, each essential in the metastatic cascade, were deregulated in clones able to generate brain metastases compared to others. To identify the set of genes that are most likely to be predictive of brain metastases, we took advantage of the compatibility of LeGOTek system with machine learning classification, using clonal and single cell information from the scRNA sequencing. Interestingly, we identified a gene expression signature, including – but not exclusively – genes from the neurogenesis pathways, predictive of brain tropism in the PDX dataset. As the PDX dataset only included one PDX with no brain metastasis, it will be important to validate these results in larger PDX datasets and patient samples. Further experiments will be necessary to refine this signature to a smaller number of commonly deregulated genes and study the role of individual genes in tumour progression and brain tropism.

While our results suggest that the upregulation of neurogenesis correlates with brain metastasis, it is also likely to be associated with aggressiveness in primary sites and other distant sites such as the lungs and liver. Nevertheless, other mechanisms or genes might be at play. For instance, PDX-1432, which was able to outcompete other PDXs in the primary site, showed a low score for the neurogenesis pathway and in our brain metastatic potential classifier. It will be interesting to study differences between clones showing different levels of aggressiveness, such as 4 and 6/16 for instance. While there is a risk that our phenotype of interest may be confounded by a more general aggression phenotype, the discrepancy in aggression (as quantified by growth in primary tumours and time to metastasis *in vivo*) between clones 6, 16 and 4 could provide further insights.

In summary, the use of our novel LeGO-Teknik library revealed that those breast cancer cells at primary sites with transcriptomic features akin to nerves are most likely to proficiently metastasize as clonal populations to the brain, as a consequence of organ mimicry. This study provides new insights into the mechanisms associated with clonal selection in brain metastases. Furthermore, it demonstrates that studying differences between clones within a tumour (i.e. intra-tumoral heterogeneity) can be used to inform and predict differences between tumours (inter-tumoral heterogeneity). Applying this technology and strategy to multiple barcoded models, including syngeneic models, will enable the identification of a plethora of mechanisms involved in clonal fitness and growth in specific TMEs such as the brain, to improve the management of patients with breast cancer.

## Methods

### Cloning of the LeGO-Teknik plasmids

The Teknik cassette was inserted in the original LeGO plasmids (see below) using the In-Fusion cloning kit (#638952; TaKaRa Bio) following the manufacturer’s recommendations. For each LeGO plasmid, a 52bp sequence was inserted 1196bp after the fluorescent protein putative stop codon. The inserted sequence contains a unique 30bp Teknik barcode and the common 10X Genomics Capture sequence 2 (CS2). After the cloning, sanger sequencing was performed for each plasmid to confirm the presence of the Teknik cassette. The details of the primers used, and the specific sequences are in the following table (F: forward; R: reverse):

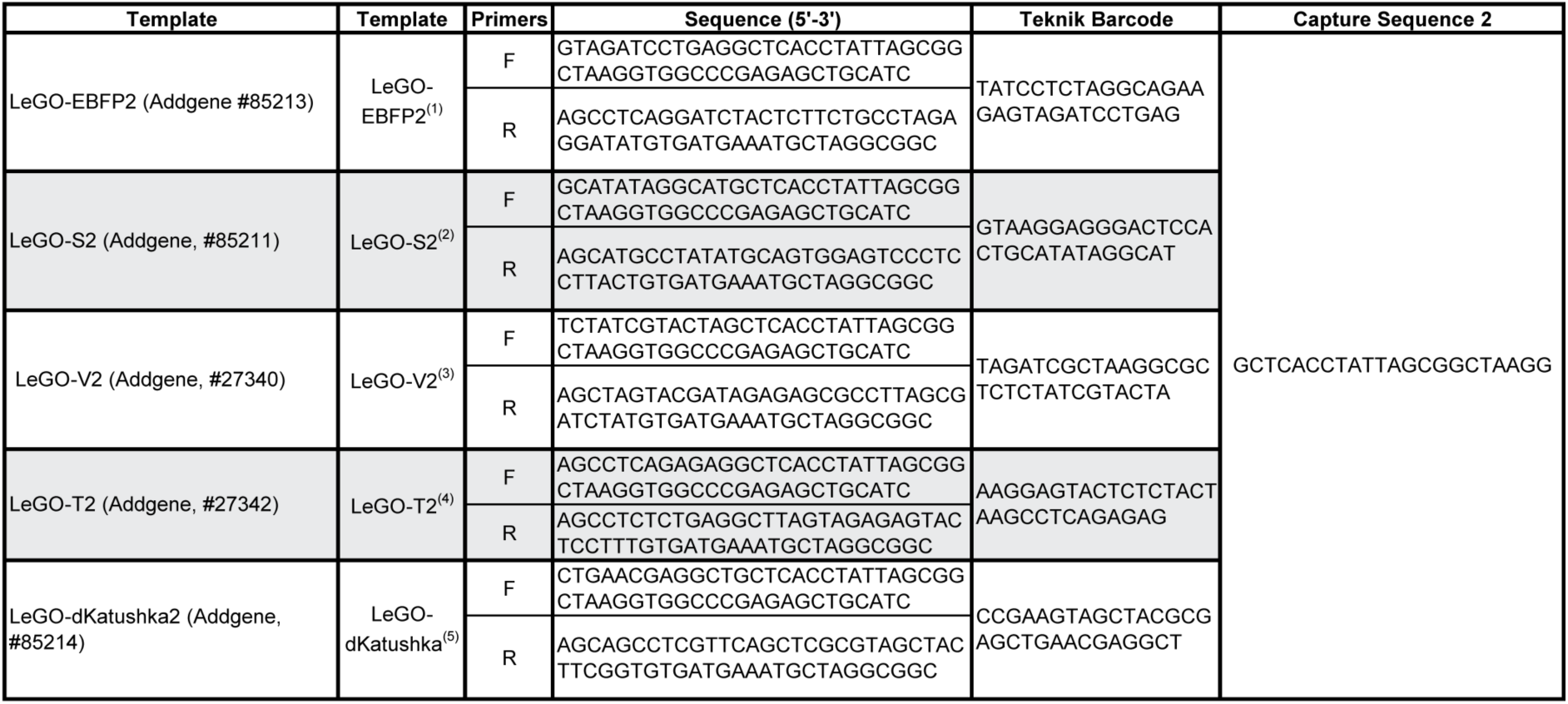

Plasmids ^(1)^ LeGO-EBFP2 (Addgene plasmid # 85213 ; http://n2t.net/addgene:85213 ; RRID:Addgene_85213);^(2)^ LeGO-S2 (Addgene plasmid # 85211 ; http://n2t.net/addgene:85211 ; RRID:Addgene_85211); ^(3)^ LeGO-V2 (Addgene plasmid # 27340 ; http://n2t.net/addgene:27340 ; RRID:Addgene_27340); ^(4)^ LeGO-T2 (Addgene plasmid # 27342 ; http://n2t.net/addgene:27342 ; RRID:Addgene_27342) and ^(5)^ LeGO-dKatushka2 (Addgene plasmid # 85214 ; http://n2t.net/addgene:85214 ; RRID:Addgene_85214) were a gift from Boris Fehse ^12,15^.

### Lentivirus production, purification, and titration

Lentiviruses were produced by transient co-transfection of human embryonic kidney–293T cells with the LeGO-Teknik vectors of interest and the packaging plasmids (pCMVR8.74, Addgene, #22036; pMD2.G, Addgene, #12259), using FuGENE HD transfection reagent (#E2311, Promega) in 10cm cell culture dishes, following the manufacturer’s recommendations. The transfection and virus production were performed in serum-free Opti-MEM (#31985070, Thermo Fisher Scientific). The DNA and FuGENE reagent were washed 4 hours after the transfection and fresh media was added to the plate. After 48 hours, the supernatant of the cells was collected and centrifuged (500g, 5min, 4°C) and the supernatant was further concentrated using an Amicon Ultra-15 100-kDa centrifugal filter unit (#UFC910024, Merck). The virus titer was then measured using the fluorescence titrating assay and the formula TU/ml = (Number of cells transduced × % fluorescent)/(Virus volume in ml).

### Cell lines

The human triple negative breast cancer cell line MDA-MB-231 was purchased from the American Type Culture Collection (ATCC; #HTB-26, passage 39). Both the MDA-MB-231 and human embryonic kidney-293T cells were maintained in DMEM high glucose (#11965092, Thermo Fisher Scientific) supplemented with 10% fetal bovine serum (FBS) (Moregate Biotech) and penicillin-streptomycin (10,000 U/ml) (#15140122, Thermo Fisher Scientific) at 37°C in a humidified atmosphere with 5% CO_2_.

### Flow cytometry

All samples were sorted using BD FACSAria III and samples were analysed using BD FACSAria III or BD FACSymphony A3, with the following configurations:

-BD FACSAria III cell sorter with BD FACSDiva 8.0.2 software (Becton Dickinson and Company, BD Biosciences, San Jose, CA, USA), equipped with four lasers and capable of detecting up to 17 parameters. Samples were analysed using the following excitation lasers and emission filters: eBFP2 (405 nm; 450/40), tSapphire (405 nm; 510/50), Venus (488 nm; 530/30), DRAQ 7 (633 nm; 710/50), tdTomato (561 nm; 582/15), and Katushka (561 nm; 670/14). For sorting, a 100-μm nozzle tip was used with a sheath pressure of 138 kPa (20PSI) and a drop drive frequency of 30 kHz. BD FACSFlow (BD Biosciences, San Jose, CA, USA) sheath fluid was used. The sample and collection tubes were maintained at 4°C.
-BD FACSymphony A3 cell analyser with BD FACSDiva 9.3.1 software (Becton Dickinson and Company, BD Biosciences, San Jose, CA, USA), equipped with five lasers and capable of detecting up to 30 parameters. Samples were analysed using the following excitation lasers and emission filters: eBFP2 (405 nm; 431/28), tSapphire (405 nm; 525/50), Venus (488 nm; 515/20), tdTomato (561 nm; 586/15), and Katushka (561 nm; 670/30), DRAQ 7 (633 nm; 710/50).

### Establishment of the LeGO-Teknik labelled MDA-MB-231 population and single cell derived clones

MDA-MB-231 cells were infected with a mixture of the normalised five LeGO-Teknik lentiviruses. The combination of the five different fluorophores resulted in 31 different colours. The volume of viruses needed to reach 75% positive cells was empirically determined. The cells were washed and sorted by flow cytometry, 72 hours after the infection. For each colour, 100 cells were sorted, and the mixture of the 31 colour-coded populations (3,100 cells in total) was plated in a 48-well plate. This normalised population was then expanded in vitro. Once a T75 flask was confluent, the cells were detached, centrifuged, and resuspended in PBS containing 2% of FCS before sort. For each colour, individual single cells were sorted directly into 48 wells of a 96 wells plate. After expansion, individual clones were analysed by flow cytometry to confirm their optical barcode and purity. Following this protocol, we were able to generate 13 single cell derived clone of MDA-MB-321 cells with a unique optical barcode. These clones were individually expanded and frozen, enabling different combinations to be mixed prior to *in vivo* injections.

### DNA extraction and library preparation

DNA extraction from 1M cell pellets was performed using the Monarch® HMW DNA Extraction Kit for Cells and Blood (NEB T3050S) following the manufacturer’s protocol, setting the lysis to 600 rpm. Eluted DNA was homogenised and sheared to a 30 kb target size using 10 passes through a 29G needle. The sheared DNA was quantified using the Qubit dsDNA BR Kit and analysed on the 4200 TapeStation System (Agilent) with Genomic DNA ScreenTape to confirm proper fragmentation and concentration. The Ligation Sequencing Kit (ONT SQK-LSK114) was used for whole genome sequencing preparation and library loading on PromethION R10.4.1 flow cells (PRO0114M, one flow cell per clone), following the Oxford Nanopore Technologies protocol “Human variation sequencing from 30 kb extracted cell line samples using SQK-LSK114.” Sequencing was performed on a P2 solo system, including 2 washes and reloads per flow cell.

### Genomic data analysis

Dorado v0.9.5 was used to basecall the reads offline in “sup” accuracy (dna_r10.4.1_e8.2_400bps_sup@v5.0.0), including modified mCG and hmCG (5mCG_5hmCG v3), and align to a human reference genome hg38 (GCA_000001405.15_GRCh38_no_alt_analysis_set.fna) concatenated with the sequences of the fluorescent tags, to generate BAM files for each clone. SAMtools version 1.21 ^34^ was employed for sorting and indexing the BAM files.

The human variation workflow (wf-human-variation) from EPI2ME was used for calling single nucleotide variants (SNVs), structural variants (SVs), and copy number variants (CNVs), and phase the alignment files. Phylogenetic relationships between clones were determined from the SNV data as follows: individual vcf files were merged using bcftools merge ^34^, and well supported variants with a quality score QUAL > 20 and read depth DP > 10 in at least one of the clones were retained. For overlap calculations, a SNV was considered to be present in a clone if at least 1 copy was present (thus encompassing both heterozygous and homozygous calls). Overlaps were plotted using the ComplexUpset R package ^35^. The filtered multi-sample vcf file was analysed using the SNPrelate R package ^36^ for hierarchical clustering based on identity-by-state analysis.

### Establishment of the LeGO-Teknik labelled PDX cells

PDX-0066 and PDX-1432 were established at the Olivia Newton-John Cancer Research Institute (ONJCRI, Australia). PDX-1432 was generated from a drug naïve TNBC BRCA-1 wild type tumour and PDX-0066 from a malignant pleural effusion of a patient with BRCA-1 mutated TNBC. The PDX CRCM412 (PDX-412), PDX CRCM434 (PDX-434), PDX CRCM226 (PDX-226) (Cancer Research Centre of Marseille) were generated at the Institut Paoli-Calmettes (France) from drug naïve TNBC tumours (PDX-434, PDX-412) or drug naïve HER2 amplified tumour (PDX-226). The use of patient samples was approved by Austin Health Human Research Ethics Committee.

Single-cell suspensions from each PDX (passage 6 for PDX-0066, passage 10 for PDX-226, passage 10 for PDX-434, passage 6 for PDX-1432 and passage 7 for PDX-412) were prepared as described below. The cells were plated at the density of 300,000 cells per well in a 24-well flat-bottom ultralow attachment plate (#734-1584, Corning) and were kept in culture for 72 hours in 300 μl of mammosphere media containing Dulbecco’s modified Eagle’s medium F12 (#10565042, Thermo Fisher Scientific), B-27 supplement 1X (#17504001, Thermo Fisher Scientific), penicillin-streptomycin (100 U/ml) (#15140122, Thermo Fisher Scientific), insulin (5 μg/ml) (#11376497001, Sigma-Aldrich), hydrocortisone (1 μg/ml) (#H0396-100MG, Sigma-Aldrich), heparin (0.8 U/ml) (#H0878-100KcU, Sigma-Aldrich), basic fibroblast growth factor (20 ng/ml) (#01-106, Merck-Millipore), and epidermal growth factor (20 ng/ml) (#E9644, Sigma-Aldrich) in the presence of one LeGOTek virus (Teknik-eBFP2 for PDX-226, Teknik-tSapphire for PDX-0066, Teknik-Venus for PDX 434, Teknik-tdTomato for PDX 1432 and Teknik-Katushka for PDX-412). For each PDX, 100,000 positive cells were sorted and transplanted in one NSG mouse as described below. The primary tumours were harvested when they reached around 800 mm^3^. A small piece of the tumour was kept for imaging and a single cell suspension was made. The cells were viably frozen and used for further experiments, including single cell RNA sequencing and transplantation of mixed cell population.

### Overexpression of SOX4 in MDA-MB-231 clone 3

To generate SOX4-overexpressing breast cancer cells, MDA-MB-231 clone 3 cells were transduced with a lentiviral vector encoding the human *SOX4* gene (NM_003107.3) under the control of the SFFV promoter. This vector included an mCherry fluorescent marker and a puromycin resistance gene separated by a T2A self-cleaving peptide. The lentiviral vector used for SOX4 overexpression, pLV[Exp]-mCherry:T2A:Puro-SFFV>hSOX4[NM_003107.3], was obtained from VectorBuilder (Vector ID: VB250306-1458kfh). Control cells were transduced with a matching lentiviral backbone lacking *SOX4*, in which the open reading frame was replaced by a stuffer sequence corresponding to amino acids 2–83 of *E. coli* β-galactosidase. This control vector lacked the mCherry fluorescent marker but retained puromycin resistance for selection. The control vector, pLV[Exp]-Puro-SFFV>ORF_Stuffer, was also obtained from VectorBuilder (Vector ID: VB250210-1540ghb). Following transduction, cells were subjected to puromycin (#A1113803, Thermo Fisher scientific) selection (10 µg/mL) for two weeks to ensure stable integration and expression of the constructs.

### In vivo experiments

For each model, cells were injected into the exposed fat pad from the fourth mammary gland of 7-week-old NSG female mice. The cells were resuspended in 10 μl of injection buffer: 25% growth factor–reduced Matrigel (#734-1101, Corning) 42.5% DPBS (#14190144, Thermo Fisher Scientific), 30% FBS (Moregate Biotech), and 2.5% Trypan blue (Bio-Rad Laboratories) for injection visualization. Tumour volume was measured twice weekly using electronic callipers. The tumour volume was estimated using the following formula: (minimum diameter)^2^ × (maximum diameter)/2. Tumours were resected when they reached a volume of about 200 mm^3^ for the MDA-MB-231 cells and 500 mm^3^ for the different PDX models. The weight of the mice was also measured twice weekly or daily once they lost 10% of their body weight. When the weight loss reached 15%, the animals were humanely euthanized by CO_2_ excess and metastatic organs were collected for further analysis. All animal procedures were approved by the Austin Animal Ethics Committee and conducted in accordance with the National Health and Medical Research Council guidelines.

### Single-cell suspension preparation

The tissues were manually chopped into small pieces (about 1 mm by 1 mm) and resuspended in the digestion medium: collagenase IA (300 U/ml) (#C9891, Sigma-Aldrich), hyaluronidase (100 U/ml) (#H3506, Sigma-Aldrich), and deoxyribonuclease I (DNase I) (100 U/ml) (#LS002139, Worthington) in DMEM/F12 + glutamax (#10565018 Thermo Fisher Scientific) supplemented with 10% fetal bovine serum (FBS) (Moregate Biotech) and penicillin-streptomycin (10,000 U/ml) (#15140122, Thermo Fisher Scientific). The tumour was resuspended in 5 ml of digestion medium, the lungs in 5 ml of digestion medium, and the liver in 10 ml of digestion medium. Samples were incubated for 30 min at 37°C with agitation, then pass through a 70-μm cell strainer and spun down for 5 min at 500*g*. The pellet was resuspended in 5ml of TrypLE Express (#12604013, Thermo Fisher Scientific) for 5 min at 37°C with agitation. The TrypLE was inactivated by the addition of 35 ml of DPBS containing 2% FBS. The cell suspension was then filtered through a 40-μm cell strainer and spun down for 5 min at 500*g*.

### Sample preparation and confocal imaging

After harvesting, samples were immediately put in freshly made, ice-cold 4% paraformaldehyde (#28908, Thermo Fisher Scientific) in DPBS and kept in the dark, at 4°C for 16 hours. Fixed samples were then washed in DPBS and embedded in 3% low–melting point agarose (#1613111, Bio-Rad Laboratories). Sections (200 μm) were cut using a vibratome (#VT1000S, Leica Biosystems) and incubated with DRAQ5 (ab108410, Abcam) to stain the nuclei of the cells before being mounted on slides (#J1800AMNT, Thermo Fisher Scientific) with high-performance coverslips (#474030-9000-000, Carl Zeiss Microscopy) and antifade mounting medium (#P36961, Thermo Fisher Scientific). Images were acquired using LSM 980 (Carl Zeiss) confocal microscopes equipped with 20×/0.8 objectives. Online fingerprinting was used to spectrally unmix the 5 different fluorescent protein BSVTK fluorophores with reference spectra acquired in cancer cells expressing a single colour only.

### Single cell experiments using the LeGO-Teknik barcoding strategy

The experiments were performed using a Chromium controller machine (#120223, 10x Genomics) and a Chromium Single Cell 3ʹ GEM, Library & Gel Bead Kit v3 (1000075, 10x Genomics) and a Chromium Single Cell 3ʹ Feature Barcode Library Kit (1000079, 10x Genomics). For each experiment, 10,000 cells were captured in gel bead in emulsion. The protocol used was the “Chromium Single Cell 3ʹ Reagent Kits v3 with Feature Barcode technology for CRISPR Screening” user guide (Document CG000184, Rev C, February 2020) with some LeGO-Teknik specific variations. The GEM generation and barcoding, Post GEM-RT Cleanup and 3’ Gene expression library construction steps were unmodified. Only the cDNA amplification, Teknik-Feature library construction and the libraries pooling steps were adapted for the LeGO-Teknik strategy. All the details of the protocol, including the necessary extra reagents are available here: https://www.protocols.io/private/53518A25980911EF87650A58A9FEAC02.

### Bulk RNA sequencing of MDA-MB-231 clones

200 cells from cancer clones 3, 6 and 16 were sorted from in vitro cell culture or from single-cell suspensions of primary tumours using the BD FACSAria III Cell Sorter. Cells were sorted directly into the CDS sorting solution from SMART-Seq mRNA LP kit (#634752, TAKARA Bio) and the libraries were prepared according to the manufacturer’s instructions. The indexed libraries were pooled and paired-end sequencing was performed (2 x 61 cycles) on a NextSeq 1000 instrument (Illumina) using a P2 100 cycle kit.

### Bulk RNA sequencing of MDA-MB-231 clone 03 overexpressing SOX4

Total RNA was extracted from confluent cultures grown in 6-well plates using the Monarch Total RNA Miniprep Kit (T2010S, NEB) in combination with the NEBNext Poly(A) mRNA Magnetic Isolation Module (E7490S, NEB), following the manufacturer’s instructions. Samples included: clone 03 wild-type (WT), clone 03 control (transduced with ORF stuffer), clone 03 overexpressing *SOX4*, and clone 06 WT. The quality of isolated mRNA was assessed using the RNA ScreenTape system (5067-5576, Agilent), and concentrations were measured using the Qubit RNA Broad Range Assay Kit (Q10210, ThermoFisher Scientific). For each sample, 100 ng of total RNA was used to prepare libraries with the NEBNext Ultra™ II Directional RNA Library Prep Kit for Illumina (E7760S, NEB), according to the manufacturer’s protocol. Libraries were dual-indexed (E7600S, NEB), pooled, and sequenced on an Illumina NextSeq 1000 system using a P2 100-cycle kit. Sequencing was performed with the following parameters: paired-end reads (2 × 61 bp), dual indexing (Index 1 and Index 2: 8 bp each), and stranded library preparation. All conditions were analyzed in biological triplicates.

### Bulk RNAseq data processing and analysis

Bulk RNA-seq data were processed using an in-house nextflow pipeline (modules publicly available at https://github.com/DavisLaboratory/nf_modules). Quality control was performed with fastQC (v0.11.9) and MultiQC (v1.14), adapter trimming with cutadapt (v4.1), two-pass alignment with STAR (v2.7.10b), gene level quantification with featureCounts (v2.0.1), and raw expression matrices and metadata were exported as SummarizedExperiment (Bioconductor release 3.19) objects.

To identify genes upregulated by the over-expression of SOX4 in clone 3 *in vitro,* edgeR (v4.4.2) was used to test for differential expression and pathway enrichments tested using fgsea. Significant pathways (adjusted p-value < 0.01) were then analysed using vissE (v1.14.0) to cluster these based on their gene identity and derive coherent modules. A SOX4 over-expression gene signature of 300 genes was derived by taking the 150 genes up– (and down) regulated via a treat test (default fold-change threshold) with edgeR. This approach specifically tests for large fold-changes to identify genes with strong biological relevance. Testing of this gene-signature was then conducted using singscore (v1.24.0).

### scRNAseq data processing

Sequencing reads were pre-processed with 10x Genomics CellRanger v7.2.0 ^37^ using the GRCh38 reference (release 2020 from 10x Genomics) and including a feature set file to unpack our LeGO Teknik barcodes using the antibody capture option. To identify potential mouse contamination, sequencing reads were additionally processed with CellRanger with identical parameters using the concatenated human and mouse [GRCm39] reference genomes.

CellRanger output filtered count matrices were loaded as Seurat (v5.1.0) ^38^ objects in R (v4.4.1) ^39^ and pre-processed following the best-practice guidelines. Briefly, cells were filtered with quality control cutoffs (percent mitochondrial UMIs, number UMIs, number unique features; described in code), and count matrices were scaled and normalised with SCTransform. Principal Component Analyses (PCA) were calculated and used for Uniform Manifold Approximation and Projection (UMAP) dimension reduction using the first 30 principal components. Cells were clustered with FindNeighbours and FindClusters functions, and doublets were identified and removed using scDblFinder (v1.18.0) ^40^. Cell states were estimated with the CellCycleScoring function and regressed with SCTransform. Filtered quality control metrics (percent mitochondrial UMIs, number of genes, log number of UMIs) were visualised using the UMAP coordinates.

LeGO-Teknik feature counts were available under the antibody feature layer in the CellRanger output. These counts were loaded as a dataframe in R for processing with Tidyverse (v2.0.0) ^41^. We calculated the total LeGO-Teknik UMIs for each cell barcode and calculated the relative proportions of each of the five LeGO Teknik features for each cell. We visualised the distributions of both the LeGO-Teknik feature total counts and proportions for all cells and selected cutoffs for calling barcodes (described in code). Lastly, we translated the combinations of the five LeGO-Teknik features into the 31 possible barcodes for each cell and added these labels to the gene expression Seurat objects.

To identify and remove any mouse cell contamination, the CellRanger output raw count matrix using the concatenated human and mouse reference genomes was loaded as a separate Seurat object in R, and the percent feature sets were calculated for human and mouse genes for each cell. Cells were visualised before and after applying the described preprocessing and LeGO-Teknik barcode labelling. Cell barcodes with more than 20% of UMI counts belonging to mouse genes were filtered from the original Seurat objects (using the human reference only). The number of mouse cells (approximately 12 in total) with LeGO barcodes were noted in the code.

### Downstream scRNAseq analysis

Differentially expressed genes (DEGs) were identified using the Wilcoxon rank sum test with Seurat using FindMarkers. Genes were ranked by log2 fold change and used for gene set enrichment calculation with fgsea (v1.30.0) ^42^ and the Gene Ontology (GO) Biological Processes (BP) database from the Molecular Signatures Database—MSigDB (v7.5.1) ^43^. Pathways with Benjamini-Hochberg adjusted P-values less than 0.05 were considered significant. Significant pathways were visualised by normalised enrichment score and pathway size with ggplot2 (v3.5.1) ^44^.

Gene signature scores were calculated for each cell from DEG sets or from DEG subsets of enriched GO BP pathways. DEGs with Bonferonni corrected p-values less than 0.05 were split into up or down regulated gene lists. Gene signatures were then calculated with singscore (v1.24.0) ^45^ and visualised on UMAP embeddings.

To estimate transcription factor activity in the scRNA-seq data, SCENIC was run using the pySCENIC (v0.12.1) implementation in a custom nextflow pipeline (http://github.com/samleenz/nf_scenic). The v10 genome database and TF to motifs annotation with a 10Kb uppstream/downstream region were accessed from cisTargetDB. findMarkers from scran was used to identify clone specific regulons by testing AUCell scores. To identify regulons that were deregulated in clones 4, 6, and 16 compared to all other clones limma (v3.62.2) was used to test clone-pseudobulked AUCell scores.

To test identification of the original LeGO tags via single-cell sequencing, a custom CellRanger reference was created by concatenating the GrCh38 reference with the LeGO ORFs and shared 3’UTR nucleotide sequences and indexed using CellRanger mkref. The previously published ID13 dataset was re-quantified using this custom reference using CellRanger Count ^17^. The data were then read into R using DropletUtils (v 1.24.0) and the LeGO ORFs split into a separate data layer ^46^. Standard preprocessing was performed to remove low quality cells and doublets before LeGO ORF abundances were visualised on UMAP coordinates of the gene expression data. We estimate the rate of detection of clone 13 (eBFP+Venus+tdTomato+) from the number of cells with non-zero counts for all three tags, and no detected counts for the tSapphire or Katushka tags.

### Machine learning model of brain tropism in MDA-MB-231

An elastic net model was developed to classify individual cells as positive or negative for brain metastatic potential using single cell RNA sequencing gene expression data. The single-clone MDA-MB-231 data was used as clones 4, 6, and 16 were shown to be brain metastatic *in vivo* and all other clones were not observed to metastasise to the brain. To set up a binary classification task, cells from clones 4, 6, and 16 were assigned to the positive class (brain tropism) and all other clones to the negative class. This was then modelled using elastic-net regularized binomial logistic regression.

*Data preprocessing:* Cells without a confident clone annotation were removed and counts normalised with scuttle (v1.10.2). A training cohort was defined by randomly sampling 80% of the cells, with the remaining 20% held out for model validation (6,052 cells in the training cohort and 1,513 in the test cohort). Overly sparse genes (here, defined as a gene that was not detected in more than 25% of at least one clone) were excluded in the training cohort prior to model training. This was done to avoid selecting genes that may not have had a robust pattern of expression. After filtering for sparsity, 8,564 genes remained. The log-normalised abundances for these genes were then z-score transformed in the training cohort, and the per-gene means and standard-deviations retained for processing of the testing cohort. Due to the unbalanced class distribution (∼75% of cells were in the negative class) sample weights were defined as the inverse of the class proportion in the training set.

*Model fitting:* The glmnet package (v4.1-8) ^47^ was then used to select an optimal alpha value using repeated 10-fold cross validation across a sequence of alpha values (0,1]. Training set classification accuracy (measured by area under the curve at a prediction threshold of 0.5 (AUC)) was not seen to depend strongly on the selected value of alpha so an intermediate value of 0.5 was selected. The regularisation parameter, lambda, was selected to be the largest that gave a cross-validation error within 1 standard-error of the best performing lambda in the training cohort. After training, the final model consisted of 321 genes with non-zero weights.

*Model validation:* The test cohort was z-score normalized using the training cohort mean/sd and class predicted with the fitted model. Test set performance was assessed with caret (v6.0-94) ^48^. In this test set, 38 negative cells were misclassified as positive, and 23 positive cells predicted to be negative. To test the model in PDX samples, the dataset containing 5 barcoded PDX lines was used. These were re-normalised and z-scored as for the test data before the model was applied. As these PDXs were not clonally resolved, we expected to see a greater diversity of brain metastatic potential within each individual PDX than the clonal MDA-MB-231 data hence we focused on analysing the positive class prediction probabilities (a continuous score) as opposed to thresholded prediction classes.

To test gene-set over-representation in the GO biological pathways hypergeometric testing was performed using fgsea. A minimum pathway size of 50 was specified and the background set to the 8,564 genes used as input for the model.

*Alternative training scheme:* To test whether the choice of test/train split methodology had an impact on model performance, a separate training scheme was implemented where one clone at a time was used as the held-out test set. For each of the 13 iterations; gene filtering, normalisation, and calculation of class weights were repeated using the 12 remaining clones as the training set. Model fitting with the training sets and testing on the held-out set was then done as above.

## Data availability

Data will be made available at the time of publication and upon request.

## Code availability

All commands and scripts for bioinformatics analyses will be available on GitHub (https://github.com/tphlab/legoteknik_paper) at time of publication.

## Extended Data figures and captions

**Extended Data Fig. 1:**
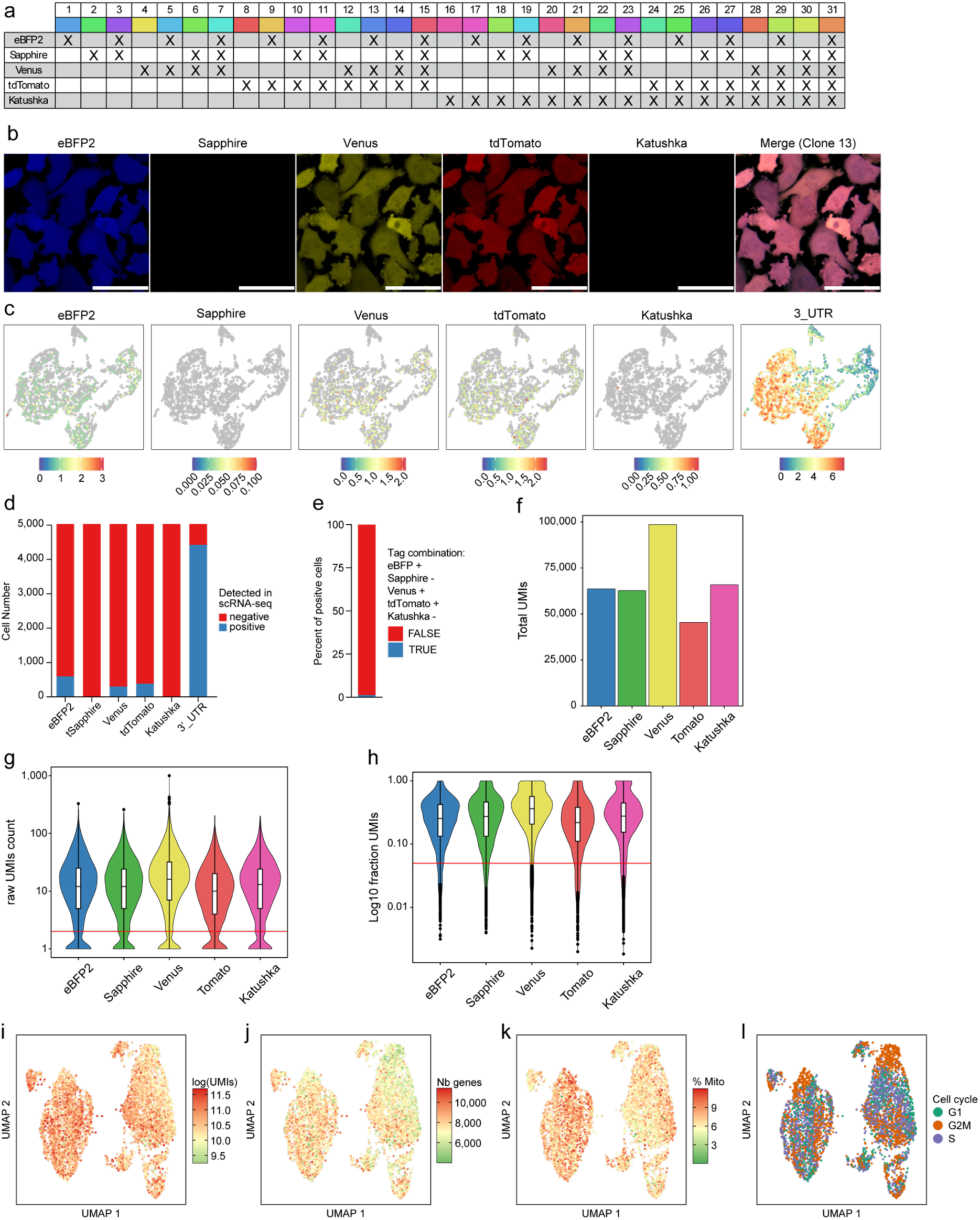
Optimisation of LeGO-Teknik optical barcoding and molecular characterisation. **a**) Multiple LeGO-Teknik colour combinations, based on the associated LeGOTek tags. The colours are conserved in all figures of the manuscript. **b**) Example of MDA-MB-231 cells ID13, expressing eBFP2, Venus and tdTomato after conventional LeGO lentivirus, generated in ^17^. The cells have been imaged by confocal microscopy (scale bar = 50µm). **c**) UMAP projection of the scRNAseq data from MDA-MB-231 ID13 cells (using conventional LeGO). The UMAP are coloured based on log2 transformed counts of the coding sequence of each fluorescent protein or the 3’ UTR region if the conventional LeGO construct. **d**) Detection rates of LeGO fluorescent proteins in chromium v3 sequencing in MDA-MB-231 ID13 cells (expressing eBFP2, Venus and tdTomato). **e**) Cellular assignment rate of ID13 with conventional LeGO tags in cells where at least one tag was detected in **c** and **d**. Cells are eBFP2/Venus/tdTomato positive and Sapphire/Katushka negative, indicating that most cells are falsely assigned using this strategy. **f**) Total UMIs from all cells for each LeGO-Teknik fluorescent barcodes. **g**) Distribution of LeGO-Teknik fluorescent barcodes showing UMI counts for each fluorescent barcode, including low cutoffs used for decoding LeGO Teknik tags. **h**) Fraction of LeGO barcode UMIs for each fluorescent barcode, including low cutoffs used for decoding LeGO Teknik tags. Single cell gene expression UMAP embeddings of MDA-MB-231 cells infected with LeGO-Teknik coloured by: log(UMIs) (**i**), number of genes (**j**), percent of UMIs from mitochondrial genes (**k**), and estimated cell cycle state (**l**).

**Extended Data Fig. 2:**
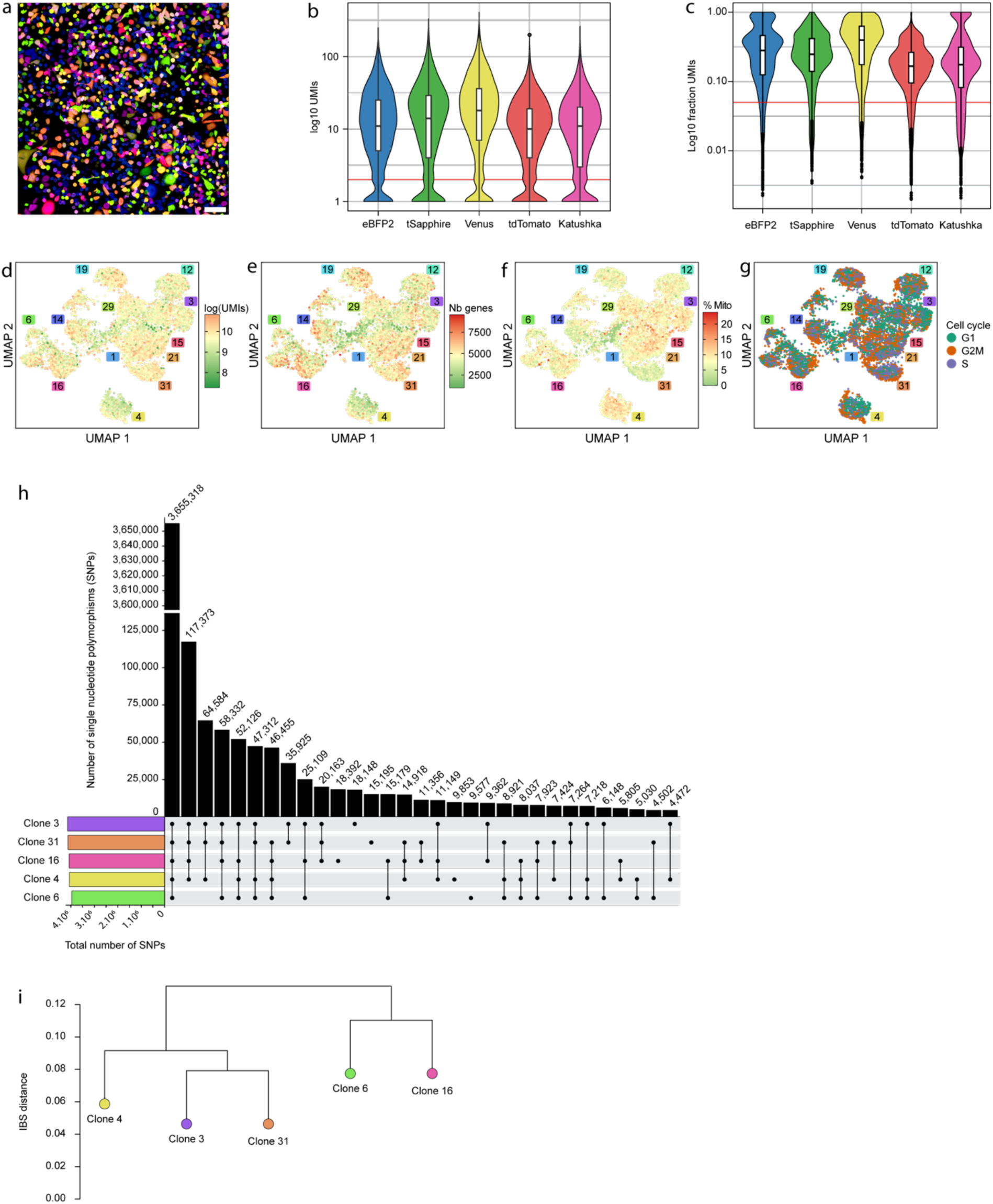
Quality control for the sequencing of MDA-MB-231 LeGOTek barcoded clones. **a**) Confocal microscopy image of 13 LeGOTek tagged single cell derived MDA-MB-231 clones in culture (scale bar = 100µm). **b**) Distribution of LeGOTek fluorescent barcodes showing UMI counts for each fluorescent barcode, including low cutoffs used for decoding LeGOTek tags. **c**) Fraction of LeGO barcode UMIs for each fluorescent barcode, including low cutoffs used for decoding LeGOTek tags. UMAP embeddings of the 13 LeGOTek tagged clones coloured by log(UMIs) (**d**), number of genes (**e**), percent mitochondrial UMIs (**f**), and predicted cell cycle state (**g**). **h**) Upset plot representing the number of SNP shared or common between the clones, determine by Nanopore sequencing of clones 3, 4, 6, 16 and 31. **i**) Phylogenetic relationships between clones based on SNP data.

**Extended Data Fig. 3:**
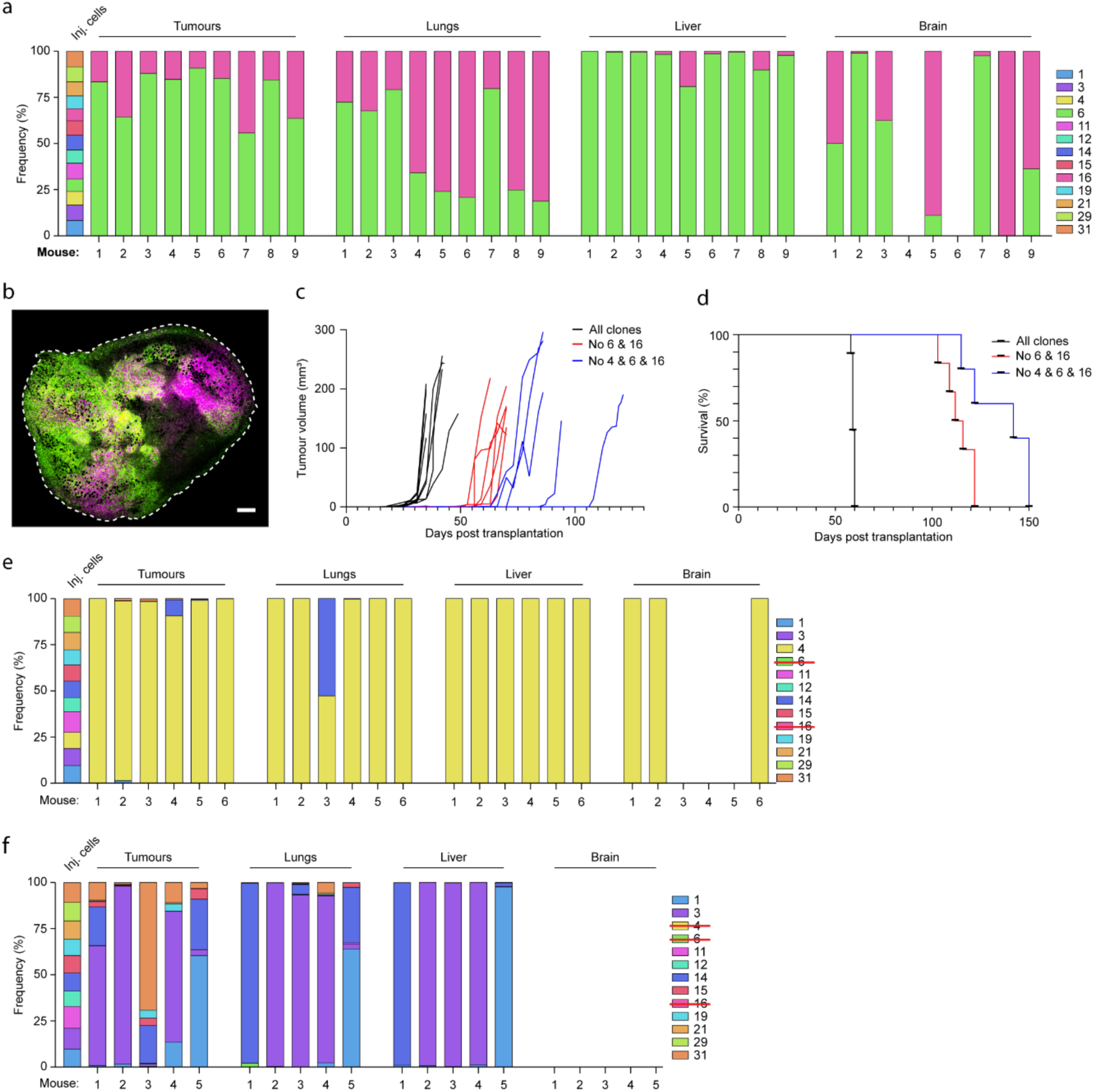
In vivo characterisation of MDA-MB-231 xenografts. **a**) Individual mice corresponding to fig 2c. Frequency of the LeGOTek tags detected in the organs of mice harvested at ethical endpoint after the tumour resection. The LeGOTek tags were detected and quantified using flow cytometry. The injected population comprised all 13 LeGOTek tagged clones. **b**) Primary tumour confocal microscopy image from a mouse injected all 13 LeGOTek tagged clones (scale bar = 500 µm). **c**) Primary tumour growth in mice injected with all 13 LeGOTek tagged clones (black line), except clones 6 and 16 (red line) or except clones 4, 6 and 16 (blue line). **d**) Survival curve for mice carrying tumour from all 13 LeGOTek tagged clones (black line), except clones 6 and 16 (red line) or except clones 4, 6 and 16 (blue line). Primary tumours were resected when they reached around 200 mm^3^ and the mice were euthanised at indicated times. **e**) Individual mice corresponding to fig 2d. Frequency of the LeGOTek tags detected in the organs of mice harvested at ethical endpoint after the tumour resection. The LeGOTek tags were detected and quantified using flow cytometry. The primary tumour was resulting from the injection of all 13 LeGOTek tagged clones except for clones 6 and 16. **f**) Individual mouse corresponding to fig 2e. Frequency of the LeGOTek tags detected in the organs of mice harvested at ethical endpoint after the tumour resection. The LeGOTek tags were detected and quantified using flow cytometry. The primary tumour was resulting from the injection of all 13 LeGOTek tagged clones except for clones 4, 6 and 16.

**Extended Data Fig. 4:**
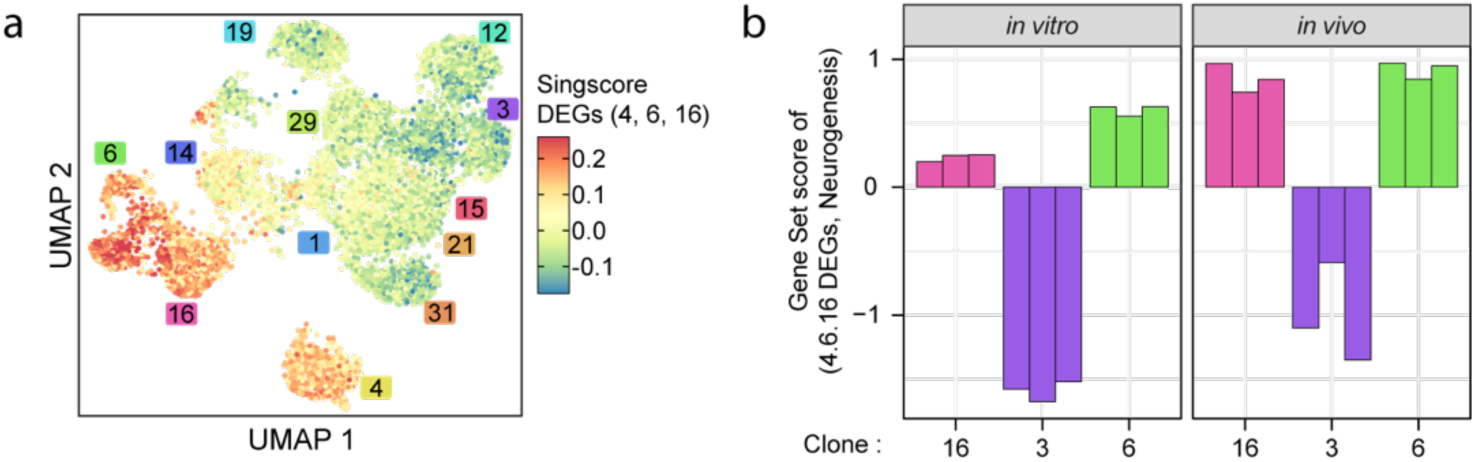
Expression of DEGs in vitro and in vivo. **a**) UMAP embeddings of the 13 LeGO-Teknik tagged clones coloured by Singscore score for the differentially expressed genes between clones 4, 6, and 16 compared to others. **b**) Singscore scores (z-score normalised) of the single cell MDA-MB-231 4/6/16 DE gene signature on bulk transcriptomes of clones 3, 6, and 16 after 200 cells sorting from cell culture (left panel) or mouse primary tumours (right panel).

**Extended Data Fig. 5:**
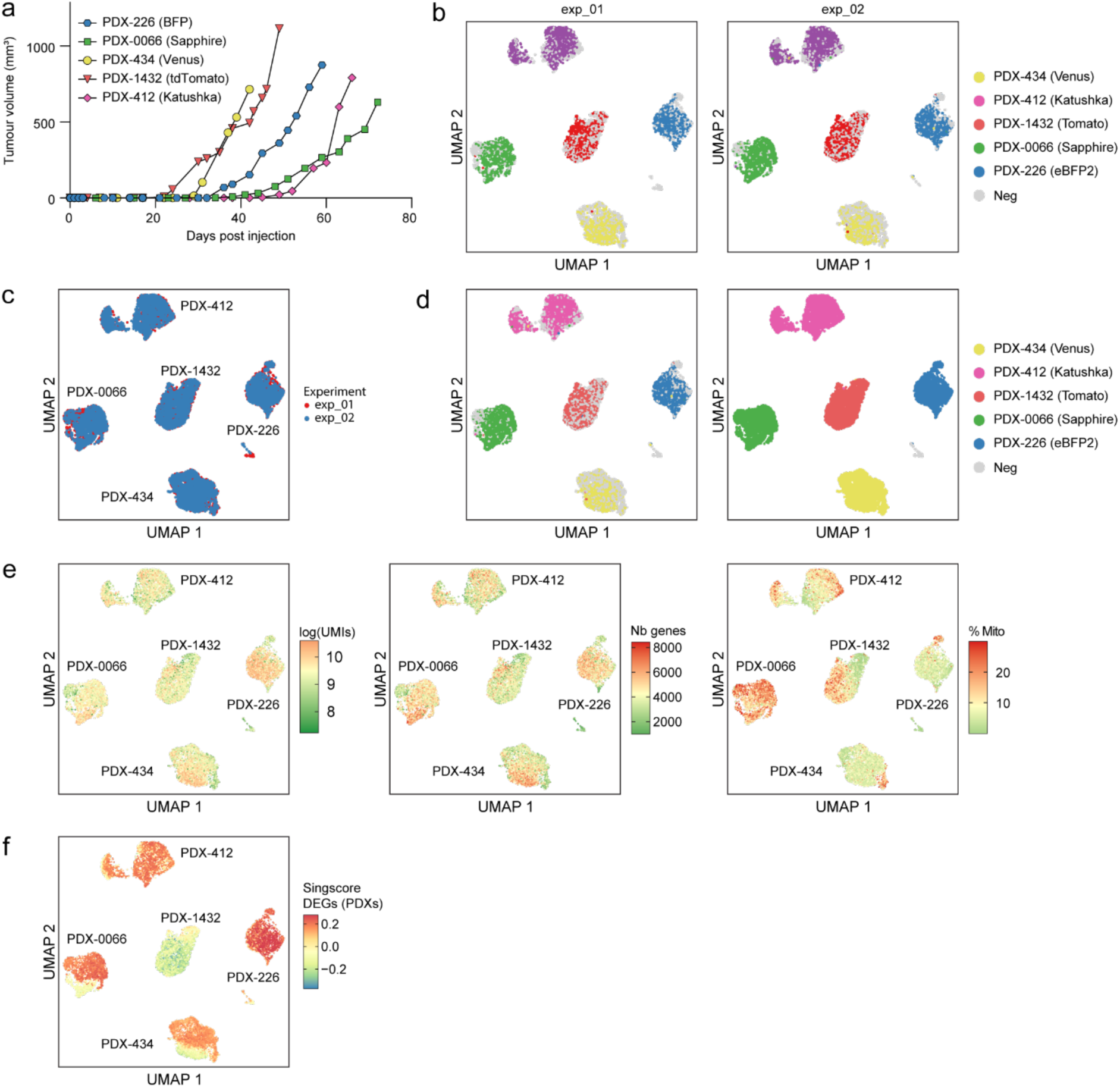
Tumour growth and single cell RNA sequencing analysis of Patient-Derived Xenografts. **a**) Tumour growth rate following injection of individual PDX in mice for the initial barcoded PDX amplification. **b**) UMAP embeddings of single cell gene expression of 5 PDXs, coloured by decoded LeGOTek barcodes for sequencing replicate exp_01 (*left*) and exp_02 (*right*). **c**) Merged UMAP embeddings from b coloured by sequencing experiment. **d**) Single cell-based-(left) and cluster-based (right) LeGOTek tag annotations. **e**) UMAP embeddings of single cell gene expression of 5 PDXs, coloured by: log(UMIs) (left), number of genes (middle) and percent of UMIs from mitochondrial genes (right). **f**) UMAP embeddings of single cell gene expression of 5 PDXs, coloured by Singscore scores of differentially expressed genes in PDX able to colonise the brain and PDX-1432.

**Extended Data Fig. 6:**
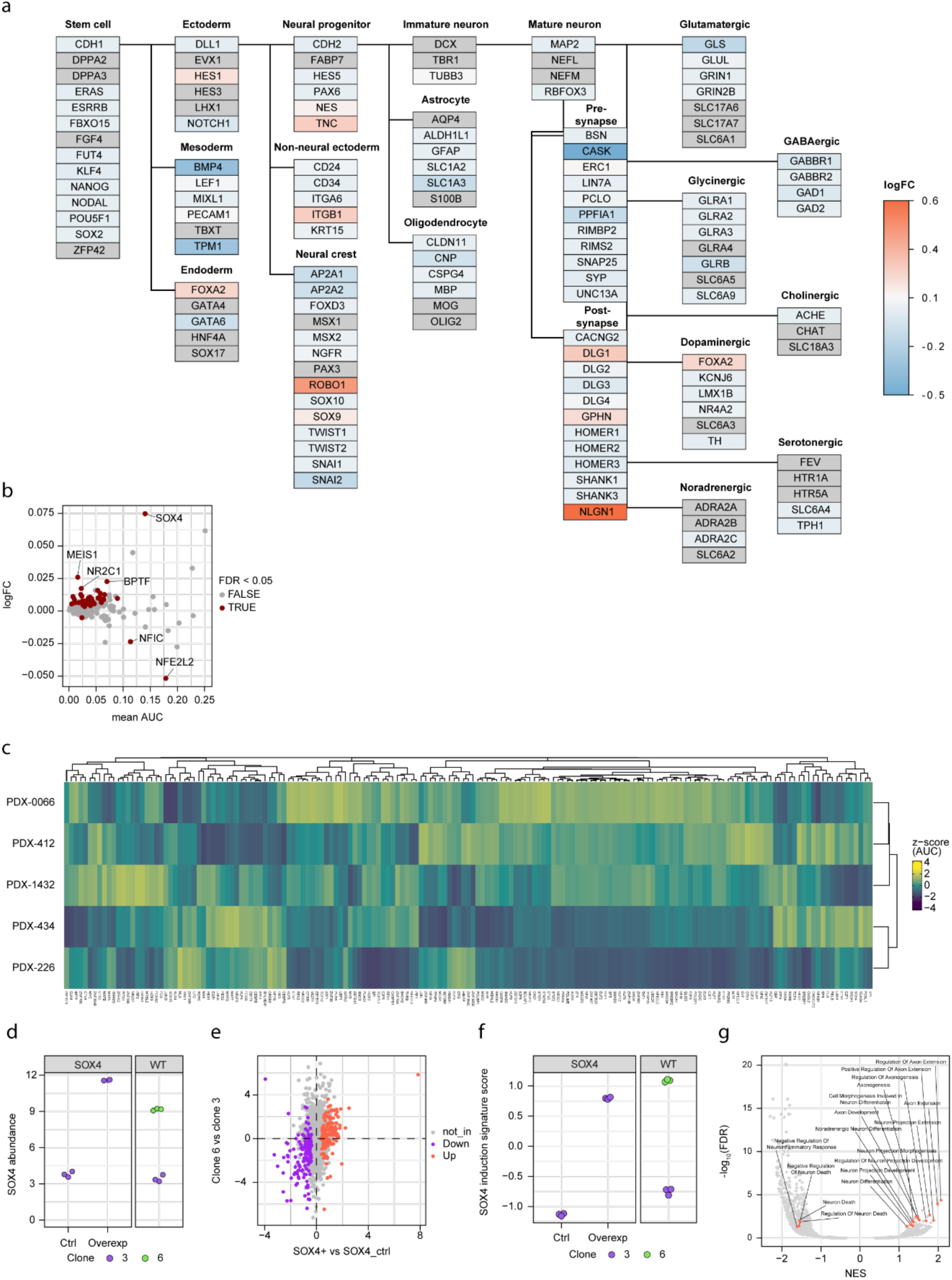
Analysis of the predictive of brain metastasis in MDA-MB-231 and PDXs. **a**) Neural lineage marker differential expression between MDA-MB-231 clones 4/6/16 and others highlights enrichment of brain metastatic clones for post-synaptic signalling lineage markers. Wikipathway ID: WP5417 (Cell lineage map for neuronal differentiation). Missing genes had their fill colour set to grey. **b**) Pseudobulk limma differential abundance testing of SCENIC regulons between clones 4, 6, 16 and others. c) SCENIC AUCell scores (z-score transformed) for the MDA-MB-231 derived regulons in the PDX dataset **d**) TPM abundance of the SOX4 gene after overexpression of SOX4 (or ORF control) and in ‘wild-type’ clones 6 and 3 by bulk-RNASeq. **e**) log-fold change plot of clone 3 SOX overexpressing vs control cells versus clone 6 WT vs clone 3 WT, genes in the SOX4 overexpression signature are highlighted in orange and purple. **f**) SOX4 gene-set scores (z-score transformed across all samples) in the SOX4 overexpressing cells (left panel) and the WT clone 3 and 6 cells (right panel). **g**) Gene-set enrichment analysis of the SOX4 over-expressing cells vs control with significant (FDR < 0.05) neural associated gene-sets highlighted in text.

**Extended Data Fig. 7:**
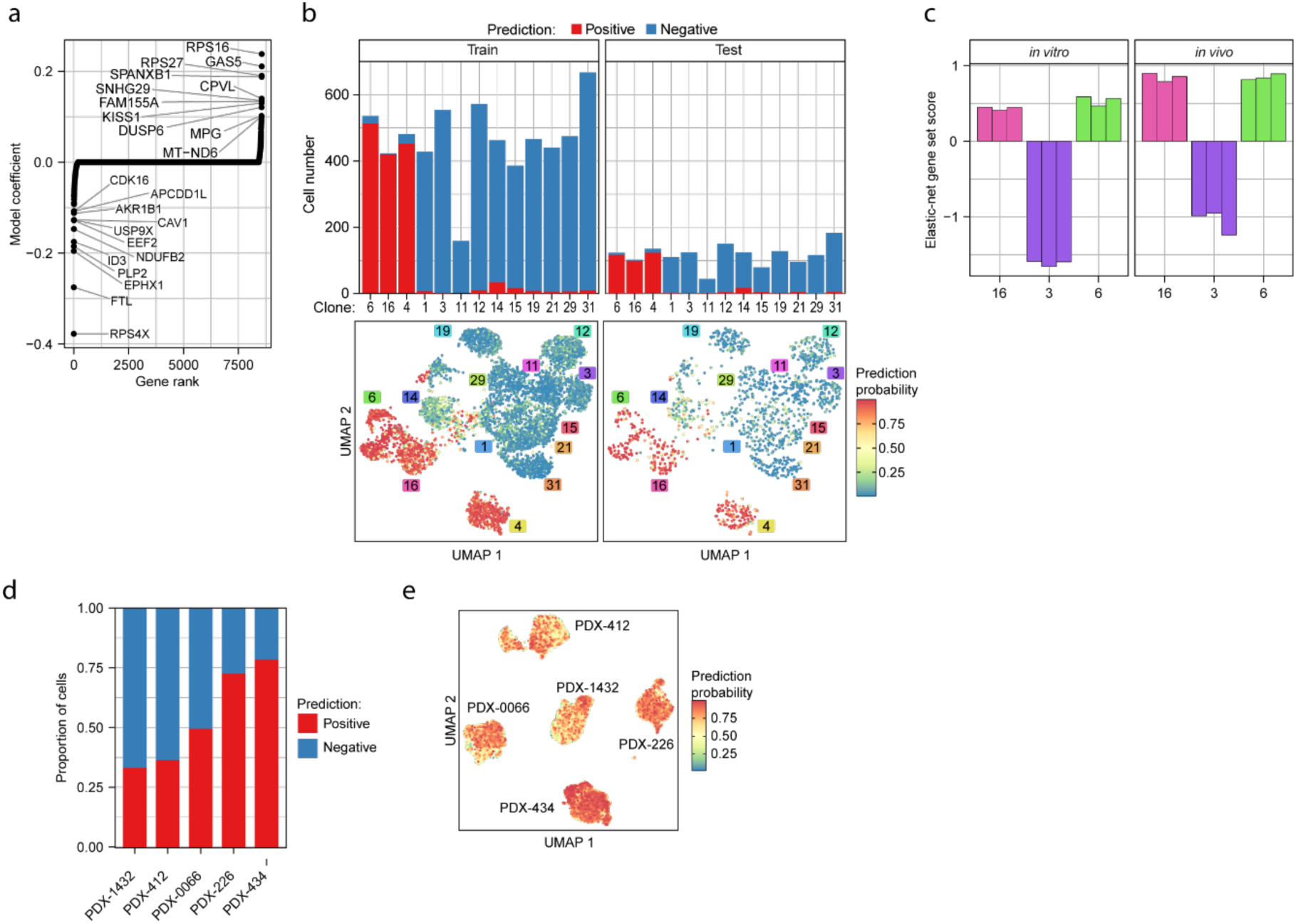
Analysis of the predictive of brain metastasis in MDA-MB-231 and PDXs. **a**) Elastic net feature coefficients in the final fitted model, ordered by magnitude of coefficient. Genes with an absolute coefficient > 0.1 are labelled. **b**) Elastic net model assignment of brain metastatic positive and negative cells in the training (top-left panel) and testing (top-right panel) cohorts in MDA-MB-231 and UMAP embeddings of MDA-MB-231 cells from the training (bottom-left panel) and testing (bottom-right panel) cohorts coloured by positive class prediction probabilities. **c**) Singscore scores (z-score normalised) of the 321 genes in the final model on bulk transcriptomes of clones 3, 6, and 16 after 200 cells sorting from cell culture (left panel) or mouse primary tumours (right panel). **d)** Elastic net model class prediction in the PDX dataset at a prediction threshold of 0.5.**e**) UMAP embedding of PDX cells coloured by positive class prediction probabilities, PDX source for each cluster is labelled.

## Acknowledgements

We thank Sarah Ellis and David Baloyan for technical assistance, and Elgene Lim, for generating PDX-1432. We are also grateful to the patients who consented for their tissue to be donated. This project was generously supported by the Love Your Sister Foundation and NHMRC (GNT2036844). DM is supported by the Victorian Cancer agency (MCRF21011), the NBCF (Investigator Initiated Research Grant IIRS0049) and NHMRC (GNT2027459). JB and MR are supported by NHMRC (GNT2036844). This research is supported by the Victorian Government through the Victorian Cancer Agency and the Operational Infrastructure Support Program. The authors and Olivia Newton-John Cancer Research Institute gratefully acknowledge the generous support of the Love Your Sister Foundation. The authors acknowledge the ACRF Centre for Imaging the Tumour Environment at the Olivia Newton-John Cancer Research Institute for providing microscopy support. We also thank the BRF and Austin Pathology. The Novo Nordisk Foundation Center for Stem Cell Medicine, reNEW, is supported by a Novo Nordisk Foundation grant number NNF21CC0073729.

## Author contributions

J.B. led the experiments, cloned the vectors and performed most of the experiments. M.R. and S.C.L. led the computational analysis, S.C.L. developed the predictive model and did the SCENIC analysis, S.G. contributed to the sequencing experiments, S.G., F.E-S., Y.W. and C.B. helped with in vivo experiments, S.L. and Q.G. performed the Nanopore sequencing and analysed the results, N.L., J.P., A.V-P. and F.J.R assisted with bioinformatics analysis, E.C-J. and C-G. provided several patient derived xenografts, M.E. and B.P. provided some reagents and advice in research design, B.Y., L.M. and D.M conceived, supervised, led and funded the study.

